# Multiple epigenetic layers accompany the spatial distribution of ribosomal genes in Arabidopsis

**DOI:** 10.1101/2020.06.17.156299

**Authors:** Konstantin O. Kutashev, Michal Franek, Klev Diamanti, Jan Komorowski, Marie Olšinová, Martina Dvořáčková

**Affiliations:** Mendel Centre for Plant Genomics and Proteomics, CEITEC, Masaryk University, Kamenice 5, 62500, Brno, Czech Republic; Laboratory of Functional Genomics and Proteomics, National Centre for Biomolecular Research, Faculty of Science, Masaryk University, Kotlářská 2, 61137, Brno, Czech Republic; Department of Cell and Molecular Biology, Uppsala University, Uppsala 751 24, Sweden; Department of Immunology, Genetics and Pathology, Uppsala University, Uppsala 751 08, Sweden; Institute of Computer Science, Polish Academy of Sciences, Warsaw 012-48, Poland; BioCEV Imaging Methods Core Facility, Průmyslová 595, 252 50 Vestec, Czech Republic

**Keywords:** rDNA, nucleolar foci, histone variants, ChIP, super-resolution microscopy

## Abstract

45S ribosomal genes in A. thaliana (rDNA) are located in tandem arrays on termini of chromosomes 2 and 4 (NOR2 and NOR4) and encode rRNA, crucial structural elements of the ribosome. Inactive rDNA genes accumulate in the condensed chromocenters in the nucleus and at the nucleolar periphery, while nucleolus delimits the active genes. We show that a subset of nucleolar rDNA assembles into condensed foci marked by H3.1 and H3.3 histones and that progressive rDNA condensation is connected with rDNA transcriptional activity, cell ploidy and rDNA copy number. Interestingly, some nucleolar foci are reminiscent of perinucleolar chromocenters, containing the NOR4 region. We further demonstrate that rDNA promoter is a key regulatory region of the rDNA repeat and describe large involvement of repressive epigenetic mark H3K9me2 and H2A.W histone variant in rDNA activity regulation. In addition, we found euchromatic H3.3 histone enrichment at the rDNA transcription start site in actively dividing tissues, despite its accumulation in nucleolar foci containing condensed rDNA repeats.

## Introduction

Ribosomal rRNA genes (rDNA, 45S rDNA) belong to the group of essential house-keeping genes, encoding key components of the ribosome −18S, 5.8S and 25S rRNA - required for protein production. Ribosomal genes in *A. thaliana* are organized into large clusters located at chromosome termini, containing from several hundreds to thousands of rDNA repeats. Such regions represent challenging targets for investigation. In *A. thaliana* two rDNA loci (3-5 MBp each) can be found at the short arms of chromosomes 2 and 4 (referred to as NOR2 and NOR4) where 500-700 copies of the 10 kb long rDNA repeat occur (Copenhaver et al, 1995; Pruitt & Meyerowitz, 1986). Individual NORs differ in their transcriptional activity: in Col-0 ecotype, NOR2 genes are inactive and show nucleoplasmic localisation (Borysko & Bang, 1951; Pontvianne et al, 2013; Shaw & Jordan, 1995), while rDNA at NOR4 is transcribed and associates to the nucleolus as one of the nucleolus associated domains (Chandrasekhara et al, 2016). The separation of the two NORs is incomplete: the rDNA bearing chromosomes tend to fuse more often than other genomic loci (Berr & Schubert, 2007), complicating the analysis of individual rDNA fractions. In addition, transcribed rDNA genes at NOR4 represent a minority of rDNA, occurring in a largely decondensed and fragile form, even escaping detection (Huang et al, 2008). Regardless of the numerous attempts to find suitable methods to distinguish between active and inactive rDNA units (e.g. determination of 3’external transcribed spacer (ETS) variants by PCR or back crossing) (Chandrasekhara et al, 2016; Pontvianne et al, 2010), microscopy remains the method of choice. It is because active rDNA of NOR4 is spatially delimited by the nucleolus, emerging as a consequence of rDNA transcription by RNA polymerase I and represents one of the most prominent nuclear structures in plant nuclei (McKeown & Shaw, 2009; Stepinski, 2014).

Nucleolus is a protein-dense, membrane-less organelle, containing predominantly components of the rDNA transcription machinery, factors involved in nascent rRNA processing and ribosomal proteins (Montacie et al, 2017; Pendle et al, 2005). The tripartite ultrastructure of the nucleolus has been elucidated by electron microscopy (EM) using differential contrasting methods (de Carcer & Medina, 1999; Medina et al, 2000). Individual conserved domains facilitating the spatial separation of rDNA processing comprise fibrillar centers (FC), dense fibrillar components (DFC) and granular components (GC) (Lafontaine, 1958; Perry, 1966). The transcription arises at the border of FC and DFC while nascent rRNA is processed in the DFC and GC (McKeown & Shaw, 2009; Melcak et al, 1996; Smirnov et al, 2014). Two types of FCs were described so far in plant meristematic cells: heterogenous and homogenous (Risueno et al, 1982). Heterogenous FCs contain condensed as well as loosened rDNA chromatin, while homogenous FCs consist of decondensed rDNA available for transcription (McKeown & Shaw, 2009), suggesting another level of compartmentalisation of nucleolar rDNA fraction. In mammals, nucleolar size correlates with the RNA pol I activity, determining the level of rDNA transcription and reflecting the cell differentiation state (Derenzini et al, 1988b; Watanabe-Susaki et al, 2014). Transcriptional activation of rDNA is maintained epigenetically. In *A. thaliana*, disruption of genes encoding HISTONE DEACETYLASE 6 (HDA6), histone methyltransferases TRITHORAX-RELATED PROTEIN 5 and 6 (ATXR5, ATXR6; H3K27me deposition) or SU(VAR)3-9 HOMOLOG 5 and 6 (SUV5, SUV6; H3K9me2 deposition) shows changes in rDNA histone methylation and acetylation and deregulation of rDNA transcriptional control (Earley et al, 2010; Mohannath et al, 2016; Pontvianne et al, 2012; Probst et al, 2004). At the global level, repressive marks H3K27me1 and H3K9me2 are enriched in ribosomal genes of *A. thaliana*, reflecting the high proportion of inactive genes; the levels of H3K4me3 and H3K27me2-3 are detectable but much lower (Mathieu et al, 2005). Similarly, both H3K4me3 (active) and H3K9me2 (repressive) histone modifications regulate the rDNA epigenetic states in the hybrid of *A. thaliana* and *A. arenosa* called *A. suecica* (Lawrence et al, 2004).

Another layer of epigenetic regulation is the incorporation of H3 histone variants. Consistent with the transcriptional activity of nucleolar rDNA, transiently expressed AtH3.3-GFP formed nucleolar foci in tobacco leaves and similar sites were found in *A. thaliana* AtH3.3-FLAG-HA tagged lines (Nie et al, 2014; Shi et al, 2011). Even though H3.1 and H3.3 differ only in four amino acids (Ahmad & Henikoff, 2002; Shi et al, 2011), these minor differences play essential role during their post-translational modification and functional specification. H3.1 is incorporated during the S phase by histone chaperone CHROMATIN ASSEMBLY FACTOR 1 (CAF-1) (Campos et al, 2015; Smith & Stillman, 1989; Tagami et al, 2004) whereas H3.3 associates with actively transcribed genes and it is deposited independently of replication by other histone chaperones (Duc et al, 2017; Duc et al, 2015; Nie et al, 2014). Chromatin immunoprecipitation – sequencing (ChIP-seq) revealed that H3.1 is abundant in heterochromatin regions, transposable elements, sites containing repressive histone marks H3K9me2, H3K27me1 or H3K27me3 and genes with methylated promoter regions (Stroud et al, 2012). H3.3 enrichment, in comparison, correlated with CG methylation in gene bodies, H3K4me1, ubiquitination of H2B and its levels peak at the 3’ end of actively transcribed genes (Stroud et al, 2012). In addition, positioning of nucleosomes inside active rDNA repeats has been long discussed. Some studies suggest that this region is less associated with nucleosomes or even devoid of them compared to the inactive rDNA genes or intergenic spacers (IGS) (Zentner et al, 2011). Others demonstrated a nucleosome shift at the promoter site regulating rDNA transcription (Li et al, 2006; Zhao et al, 2016) and the presence of non-nucleosomal complexes, summarised in (Schofer & Weipoltshammer, 2018).

Our knowledge of the structure of nucleolar rDNA, its spatial separation and chromatin organisation is very limited in plants. Due to inherent technical problems with ChIP-seq analysis of repetitive genomic regions, little is known about the distribution of epigenetic marks and histone variants in active rDNA as this approach only provides an average image of mixed active and inactive rDNA (Vaquero-Sedas et al, 2012). In this paper, we combine well-established biochemical techniques with innovative microscopy approaches to study the chromatin state of rDNA copies inside the nucleolus and on its periphery. In order to reduce the complexity of rDNA chromatin, particularly of the epigenetic states coexisting in one rDNA cluster, we employed experimental lines with reduced amount of rDNA copies containing ~ 15-20% of initial rDNA amount (Pavlistova et al, 2016). We show that NOR4-derived rDNA is organised in nucleolar foci in WT, formed as a consequence of rDNA compaction and associates with both H3.1 and H3.3 histone variants. Taking advantage of a low rDNA copy plant line, we show that both nucleolar foci and perinucleolar chromocenters are the result of the folding of rDNA in plant nucleoli. We combine ChIP-seq analysis of rDNA chromatin in seedlings with microscopic, single-locus detection of the rDNA epigenetic profiles on extended chromatin fibers and show that a portion of rDNA genes in a single NOR can contain various combinations of histone modifications. Moreover, analysis of ChIP-seq data indicates the presence of a well-positioned nucleosome at the transcription start site and dynamic changes of H3 histone variants abundance at rDNA that might be key for its transcriptional regulation.

## Results

### Both histone H3 variants are found in the nucleolus and colocalise with rDNA

The nucleolar rDNA is largely decondensed but not structure-free. In proliferating root cells a network like structure containing decondensed part and condensed foci occurs, as revealed by Structured Illumination Microscopy - SIM (Dvorackova et al, 2018). We refer to these condensed foci inside of the nucleoli throughout the text as nucleolar foci (**NF**) while we call condensed foci on the nucleolar periphery perinucleolar chromocenters. The detailed chromatin organisation of this network, however, remains poorly understood. Although it was shown that HTR5-GFP (H3.3) forms nucleolar foci in differentiated cells, the situation in proliferating cells or the presence of HTR3-GFP (H3.1) in the nucleolus were not further explored (Nie et al, 2014; Shi et al, 2011). Therefore, we decided to investigate the contribution of individual H3 variants (H3.3, H3.1) to the structure of NF and to the structure of perinucleolar chromocenters containing NOR4. We used two experimental plant lines – one expressing H3.1-GFP and H3.3-RFP (referred to as RedGreen) (Otero et al, 2016) and the second containing H3.1-RFP and H3.3-GFP (referred to as GreenRed). Imaging the root nuclei of *Arabidopsis* seedlings (up to 10-d-old) revealed that both histone variants are present in the NF (RedGreen, GreenRed; Fig. 1*A*, *SI appendix*, Fig. S1*A*) as well as in perinucleolar chromocenters (Fig. 1*A* and *D*). Nucleolar signals of H3.1 and H3.3 were not identical in both tested lines. In GreenRed plants we detected the presence of smaller, monovariant foci labelled only by H3.3 (22 / 67 nuclei), while this pattern was rare in RedGreen plants (6 / 65 nuclei; Fig. 1*B*, *SI appendix,* Fig. S1*A*). This discrepancy might be caused by unbalanced levels of transgene expression, as shown previously by western blotting (Otero et al, 2016). We propose that the majority of cells contain H3.3 variant in all NF as well as in perinucleolar chromocenters. Most of these sites are also labelled by H3.1 variant and only a subset of cells shows small monovariant H3.3 labelled NF.

**Fig. 1.**
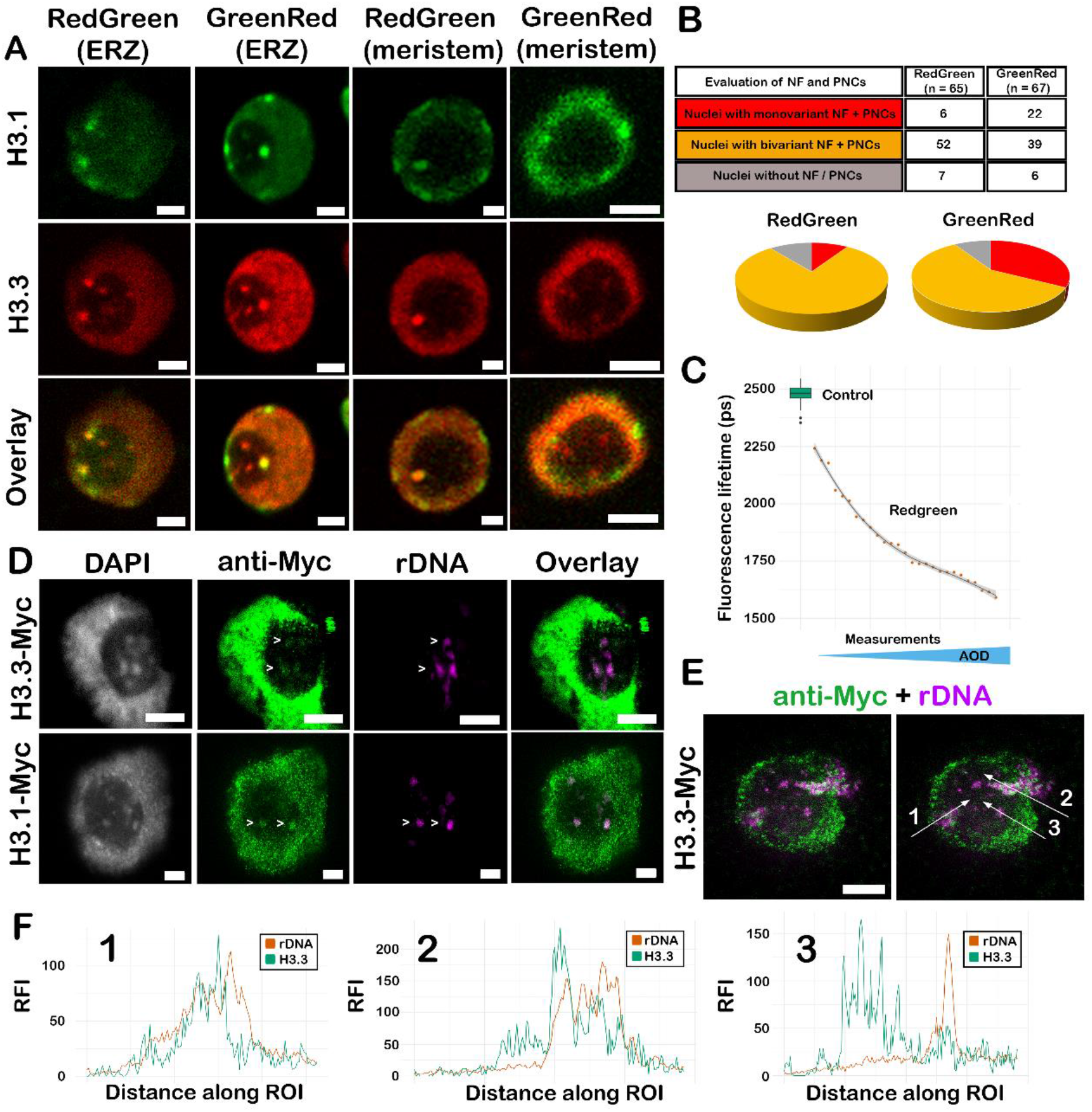
Composition of nucleolar foci (NF) and perinucleolar chromocenters (PNCs). (A) Distribution of histone H3 variants H3.1 and H3.3 in both PNCs and NF of RedGreen (H3.1-GFP; H3.3-mRFP) and GreenRed (H3.1-mRFP; H3.3-GFP) transgenic lines. Seedlings were imaged on a confocal microscope, H3.1 always color-coded green and H3.3 red for clarity. (B) Quantification of the number of monovariant (H3.3-only) and bivariant (H3.1 + H3.3) NF and PNCs observed in the nuclei of both plant lines. A minority of nuclei do not show any nucleolar foci. (C) FLIM-FRET experiment showing donor average fluorescence lifetimes (τ) for control (H3.1-GFP) and RedGreen plant lines. Box plot shows the data of the control sample. Each point belonging to RedGreen sample corresponds to a single FLIM-FRET measurement. The GFP average lifetime (y-axis) in RedGreen plants decreases with growing AOD (shown as blue triangle). (D) IF-FISH on nucleoli isolated from Myc-tagged lines showing colocalisation of H3.1 (green) and H3.3 (green) with rDNA (magenta). Images were acquired using STED microscopy (enhanced spatial resolution for green + magenta channels). (E and F) Examples of line profiles displaying various degree of overlap between H3.3 variant and rDNA (1 – strong overlap; 2 – partial overlap; 3 – no overlap in small rDNA foci). Scale bar – 2 μm. ERZ – endoreduplication zone; NF – nucleolar foci; PNCs – perinucleolar chromocenters; ROI – region of interest; AOD – acceptor over donor ratio; RFI – relative fluorescence intensity.

We further decided to investigate whether H3.1 and H3.3 are positioned in close proximity inside of the NF or form mutually exclusive environments and whether this positioning resembles the situation in the nucleoplasm. We employed Forster Resonance Energy Transfer approach combined with Fluorescence Lifetime IMaging measurements - FLIM-FRET - detecting protein-protein interactions within 10 nm. We evaluated average fluorescence lifetimes (τ) in the nucleoplasm and in the NF, using GFP and mRFP as a donor-acceptor pair. In the control experiment (H3.1-GFP plants), GFP average lifetime was τ = 2481±21.34 ps (n = 41), while in the RedGreen line (n = 27) shortening of donor lifetime was detected (Fig. 1*C*). The maximum of GFP lifetime observed in RedGreen reached τ = 2189 ps, corresponding to 12% and 36% of FRET, respectively. Since we noticed that nuclei differed in fluorophore levels, known to affect FRET (Gordon et al, 1998), we measured fluorescence intensity of H3.1-GFP and H3.3-RFP in the images acquired prior to FLIM-FRET measurement to calculate acceptor over donor (AOD) ratio. Values of this parameter varied from 20% to 86%. More pronounced lifetime shortening correlated with the growth of AOD ratio (Fig. 1*C*), explained by larger probability of energy transfer between acceptor and donor fluorophores. We repeated the experiment on the line expressing H3.3-GFP as a donor and H3.1-mRFP as an acceptor (GreenRed line). Measured FRET value was only 4.3% (*SI appendix*, Fig. S1*B*) as explained by the much higher expression of donor H3.3 than that of acceptor H3.1 in GreenRed line (Otero et al, 2016). In both GreenRed and RedGreen lines, the average lifetime of GFP-tagged histones in NF did not differ from the lifetime in nucleoplasm (*SI appendix*, Fig. S1*C*). This data suggests that these sites do not show any specific interaction or accumulation of H3.1 and H3.3 when compared to the nucleoplasm.

To test whether the NF correspond to the rDNA, we used immunodetection (IF) of individual H3 variants combined with fluorescence *in situ* hybridisation (FISH). Nucleoli-enriched samples isolated from H3.3-Myc and H3.1-Myc plant lines were imaged using STimulated Emission Depletion microscopy (STED). We observed that all H3.1 and H3.3-containing NF colocalised with condensed rDNA loci (Pearson correlation coefficient r = 0.45; SD = 0.22 and r = 0.38; SD = 0.13 for H3.1 and H3.3, respectively); (Fig. 1 *D – F*). The values of Pearson correlation coefficient were expected to be relatively low due to differences in signal intensities (rDNA signal is much stronger than that of H3 immunodetection) and enhanced resolution of STED microscopy.

### Decrease in rDNA amount leads to the loss of H3.3 labelled nucleolar foci in actively dividing root tissue

To further investigate the character of NF we employed the low-copy rDNA line 9 (L9) generated in our previous work (Pavlistova et al, 2016) and introduced a H3.3-GFP construct into this line. L9 contains only about ~15% of rDNA copies (compared to the wild-type – WT, *SI appendix*, Fig. S2) and the majority of these copies are considered active. We found that in the control WT line, H3.3-GFP forms NF, similarly to RedGreen plants (Fig. 2*A(1)*; *SI appendix*, Fig. S3 *A* and *C*). In the L9 these NF were less frequent, particularly in the meristematic root zone (Fig. 2*A(2)*; *SI appendix,* Fig. S3*B)*. When we counted the H3.3 signals in the nucleolus, we noticed a significant decrease in the number of H3.3 foci in the L9 (Fig. 2*B)*. Most of the scored from the low rDNA copy lines were observed in the elongation zone and the root tip, where the cells undergo endoreduplication and differentiate (*SI appendix*, Fig. S3 *C* and *D*). This data suggested that the formation of H3.3-labelled NF relates to the total rDNA amounts.

**Fig. 2.**
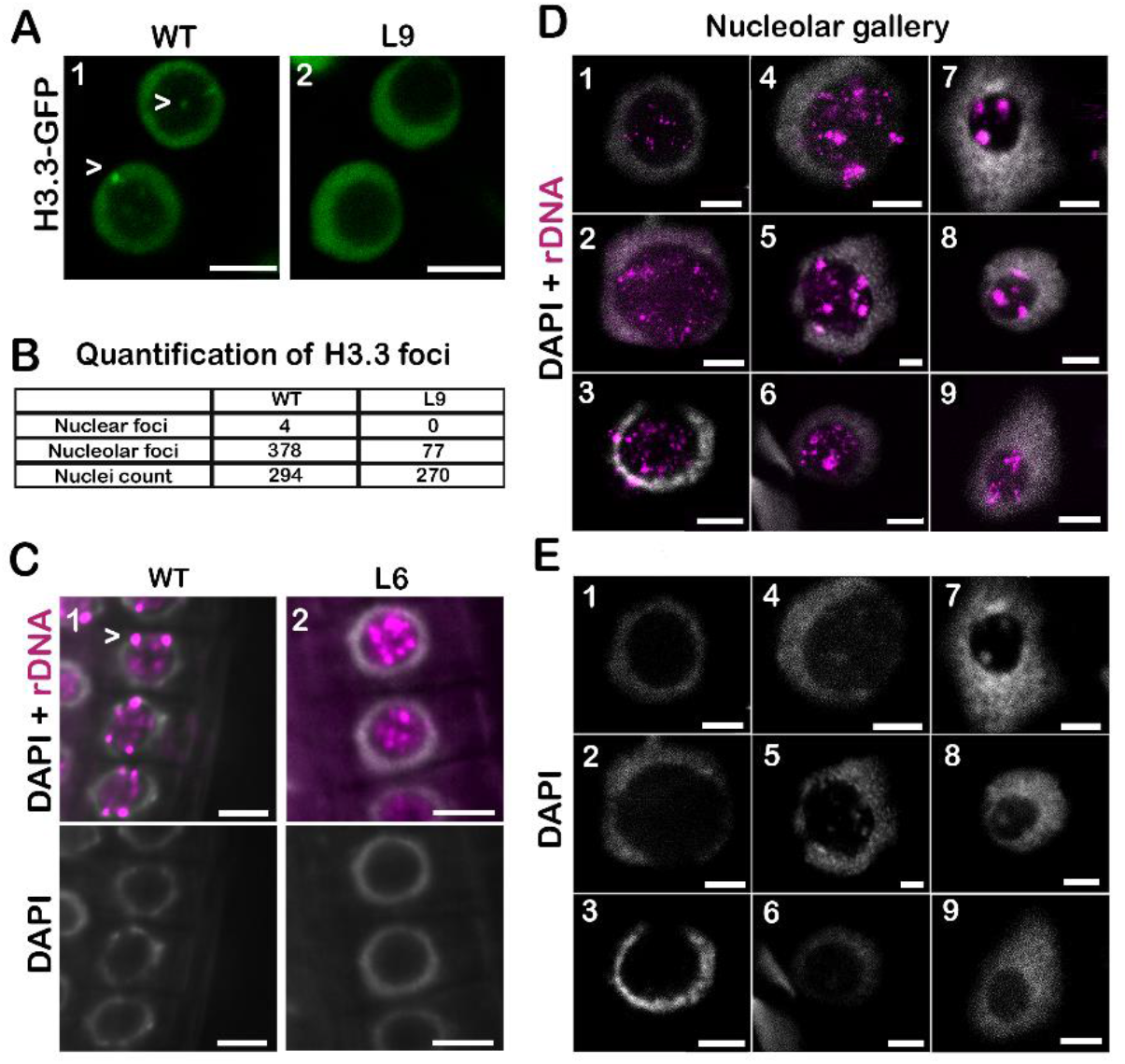
Organization of nucleolar foci (NF) and perinucleolar chromocenters (PNCs) in low-copy lines. (A and B) Analysis of histone H3.3 foci in the roots of H3.3-GFP transformed wild-type (WT) and low-copy rDNA (L9) plants by confocal microscopy. Data shown in (B) were counted from 22 images of WT roots and 17 Images of low-copy roots (3 roots per sample). (C) Organization of rDNA in the roots of WT and low-copy rDNA (L6) line using whole-mount rDNA FISH (DAPI - gray, rDNA - magenta), wide-field microscopy combined with deconvolution. (D and E) Analysis of rDNA ultrastructure using STED, rDNA (magenta), DAPI (grey, image E), merged image (D). (D1-D3) Small rDNA foci, (D4-D6) larger clusters of rDNA together with dispersed rDNA signals, (D7-D9) compact foci without dispersed rDNA signals. Scale bars – 5 μm (A and C); 2 μm (D and E).

We next performed whole-mount *in situ* hybridisation on seedlings using a rDNA probe and assessed the distribution of ribosomal genes in root cells. In WT, we detected rDNA signals both inside of the nucleolus and on the nucleolar periphery. As expected, perinucleolar rDNA signals were universally much stronger than intranucleolar ones (Fig. 2*C(1)*; *SI appendix*, Fig. S3*F*). In the low-copy line 6 (L6), where only 50-70 copies of rDNA occur in 2C cells (Pavlistova et al, 2016), perinucleolar rDNA signals were mostly absent and decondensed nucleolar rDNA sites became more pronounced (Fig. 2*C(2)*; *SI appendix, Fig. S3H)*. At this level of resolution, we could not investigate the nucleolar rDNA in more detail. Therefore, we used STED microscopy to gain a greater insight into the rDNA organization in WT and low-copy lines. The current idea of rDNA organization is based on the analysis of differentiated cells in WT plants (Pontvianne et al, 2013), where a large proportion of rDNA is inactive and which does not accurately capture the full spectrum of rDNA spatial arrangements. Improved resolution (~ 200 nm) of SIM, was recently used to examine the details of rDNA architecture in replicating cells (2C - 4C)(Dvorackova et al, 2018). STED (~ 50 nm resolution) revealed that most of the nucleoli contain NF brightly stained with DAPI and labelled by rDNA probe, resembling nuclear chromocenters. STED experiments clearly demonstrated the presence of smaller DAPI-free rDNA foci that can be found alongside larger rDNA/DAPI bright sites (Fig. *1D*). When we examined isolated nucleoli, WT and low-copy plants displayed several patterns of rDNA organization, owing to the differentiation / endoreduplication status of nuclei from which the nucleoli were isolated (Fig. 2*D*): 1) nucleoli with a big number of small, diffuse nucleolar rDNA foci occurring selectively in the low-copy line (Fig. 2 *D1* and *D2*), 2) nucleoli showing bigger and smaller foci (Fig. 2 *D3*-*D6*), 3) nucleoli with large perinucleolar chromocenters and compact nucleolar foci without dispersed smaller signals (Fig. 2 *D7*-*D9*). The latter two types of organization occur both in low-copy and WT plants. In these samples, the larger rDNA foci were DAPI labelled (Fig. 2*E*). As some nucleoli isolated from low-copy rDNA plants show similar pattern of rDNA organisation as WT (type 2 and 3), we assume that nucleolar morphology reflects the ratio of actively transcribed to total rDNA repeats. This suggests that the differences in the organization of rDNA between WT and low-copy lines are most pronounced in meristem nuclei that have the highest ratio of active rDNA copies to total rDNA content (Dvorackova et al, 2018). Consistently, the loss of the H3.3 NF occurs predominantly in the meristem root zone (*SI appendix*, Fig. S3 *A* and *B*). Once the ratio of active/inactive rDNA is low enough, the morphology of WT and low-copy nucleoli is alike (*SI appendix*, Fig. S3 *E*-*H*).

### H3.3 marks the nucleosome at rDNA transcription start site in actively dividing tissue

Since epigenetic mechanisms are essential in establishing active and inactive states of rDNA copies, we decided to investigate the distribution of histone variants and histone modifications along the rDNA repeats. We took advantage of the publicly available ChIP-seq data sets collected from young and mature leaves (actively dividing tissue with higher rDNA transcription and non-dividing tissue with lower rDNA activity, respectively) (Wollmann et al, 2017). The data were processed with an algorithm optimised for the analysis of rDNA (Zentner et al, 2011) and regions where irrelevant reads could map to (called low mappability regions, LMR, *SI appendix* Fig. S3*A*) were excluded from our analysis (described in *SI appendix, Methods*). We found that in dividing tissue, the distribution pattern as well as total levels of H3.1 and H3.3 signal were similar along the rDNA coding sequence (Fig. 3 *B* and *C*). In the non-dividing tissue, the total H3.1 level was enriched over H3.3 suggesting the silencing of a larger subset of rDNA copies typical during transcriptional downregulation. A characteristic feature of actively dividing tissues was a H3.3 peak of approx. single nucleosome width at the transcription start site, position (−73, 167) (Fig. 3*C*). This peak was lost in the non-dividing tissue, since the levels of H3.3 at the TSS dropped below 0 (Fig. 3*C*). We did not observe the accumulation of H3.3 towards the 3’end of the gene, commonly observed in RNA pol II transcribed genes (Fig. 3*D*).

**Fig. 3.**
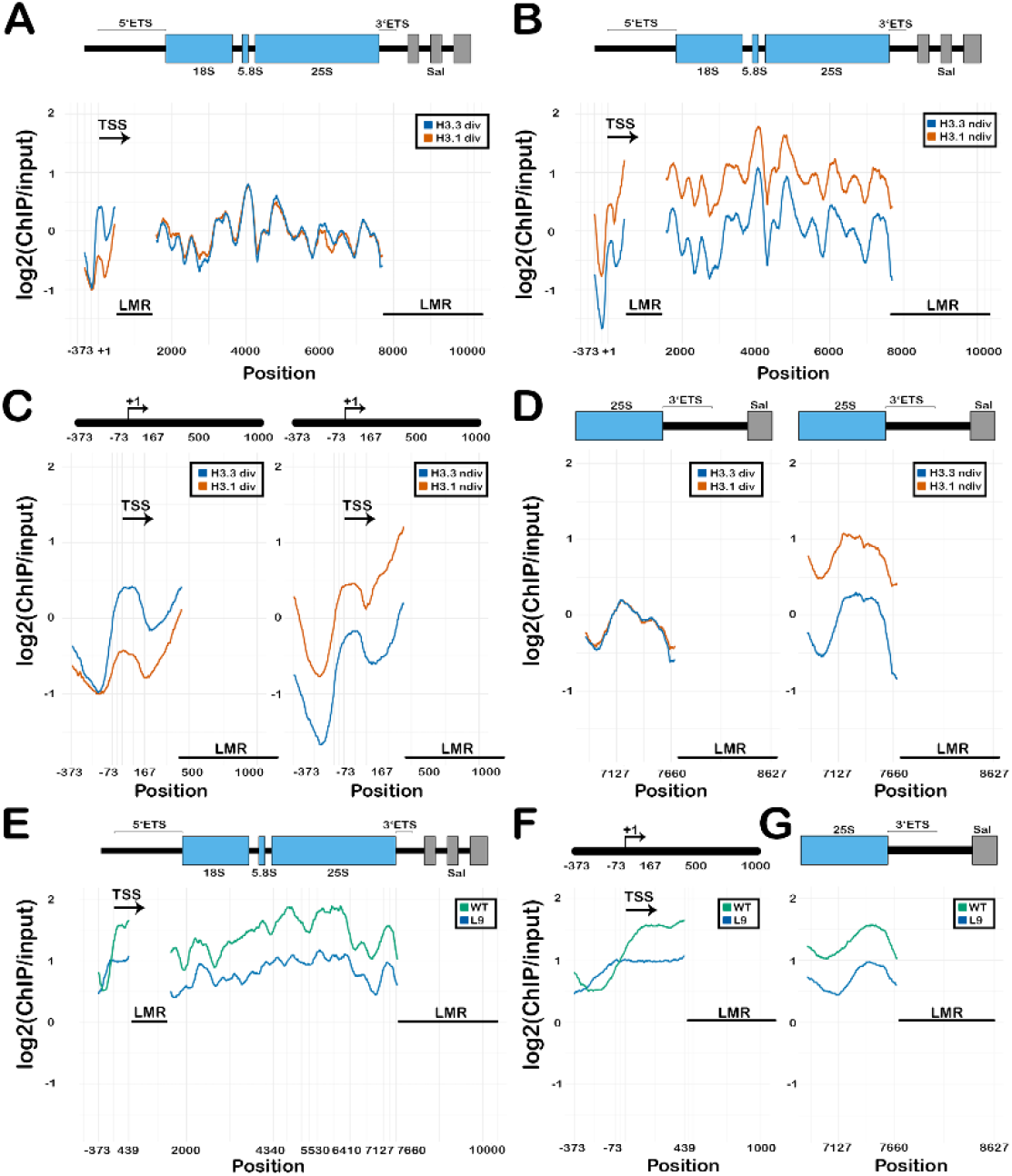
Distribution of H3.1 and H3.3 in the rDNA repeat based on ChiP-seq. (A and B) Schematic depiction of a rDNA unit used for mapping with H3.1 and H3.3 distribution profiles in (A) actively dividing (div) and (B) non-dividing (ndiv) tissues of WT plants. (C and D) Schematic depiction of mappable part of (C) rDNA promoter region and (D) rDNA TTS with profiles of H3.1 and H3.3 in dividing and non-dividing tissues (enlarged from A and B). (E) Schematic depiction of a rDNA unit with H3 histone distribution profiles in wild type (WT) and low rDNA copy line (L9). (F and G) Schematic depiction of mappable part of (F) rDNA promoter region and (G) rDNA TTS with H3 distribution profiles from WT and L9 (enlarged from (E)). ETS – external transcribed spacer; LMR – low mappability region; TSS – transcription start site.

### Wild-type and low-copy rDNA lines show similar distribution of H3 in the rDNA repeat

Since the publicly available ChIP-seq data are performed on plant lines containing WT rDNA amounts with the majority of inactive genes, we performed a ChIP-seq experiment on the low-copy line L9 (Pavlistova et al, 2016). This experiment revealed that the H3 occupancy pattern along the rDNA sequence is identical in L9 and WT (Fig. 3*E*): histone H3 is the lowest at the promoter region at the position (−323, −123) relatively to TSS followed by an abrupt signal increase at the rest of the mappable promoter region (−73, 439; Fig. 3*F*). The two peaks with the highest signal were detected at the beginning (4342, 5317) and in the middle (5317, 6427) of 25S rDNA sequence. Another feature of this profile is the increased signal level at the end of the coding sequence (7217, 7677; Fig. 3*G*).

### H3K4me3 and H3K27me3 levels are low in rDNA and mark promoter as well as the gene body

We next focused on the distribution of different histone modifications by a) using available public ChIP-seq datasets on WT and by b) employing the ChIP - Dot blot approach on the WT and two low-copy lines (L6, L9). We started with the dataset from young rosette leaves (PRJNA218138;(Chica et al, 2013)) to determine the distribution of H3K4me3 and H3K27me3 histone marks. The levels of both marks were very low in the rDNA locus, as confirmed by both Dot blot (*SI appendix*, Fig. S5 *A* and *B*) and ChIP-seq, when compared to other genomic regions (*SI appendix*, Fig. S6). The distribution pattern was similar for both modifications (Fig. 4 *A* and *B*). The lowest signal was detected at the promoter region of a single nucleosome size at (−323,−123) with respect to TSS, followed by abrupt signal increase at the remaining mappable part of the promoter adjacent to TSS (−73, 439) similarly to reported total H3 pattern. Again, largest signals were in the beginning and in the middle of 25S rDNA coding sequence at same positions as total H3 enrichment (4342, 5317; 5317, 6427) and a large peak at the end of the coding sequence, partially overlapping the H3 peak (7347, 7677). While both marks are usually associated with regulation of the promoter activity, we did not observe a significant increase in the promoter sequence in comparison with the coding sequence. This data shows very low levels of H3K4me3 and H3K27me3 in both WT and low-copy plants, suggesting a minor role in the regulation of rDNA activity.

### H3K9me2 and H2A.W are abundant in the rDNA coding sequence while H3K9me2 is lower at the promoter

The distribution of heterochromatin histone mark H3K9me2 and histone variant H2A.W was assessed using publicly available datasets (PRJNA162669; (Moissiard et al, 2012) and PRJNA219442;(Yelagandula et al, 2014), respectively). Both marks are strongly enriched in the whole rDNA locus (Fig. 4 *C* and *D*), with levels comparable to the regions of their highest enrichment such as centromeres and transposable elements (*SI appendix*, Fig. S6). Their signal is equally distributed in the rDNA coding sequence without any well-defined peaks. H3K9me2 levels are lowered by half at the promoter in comparison with the coding sequence, reaching the lowest values before the TSS (−273, −123). There is a decrease in H2A.W abundance before the TSS but it is not as pronounced and visible as with previously described modifications. Dot-blot analysis revealed a sharp decrease of H3K9me2 levels in 18S region of low-copy lines L6 and L9 when compared to WT (Fig. 4 *E*-*G*). There is less H3K9me2 in the transcription start site than in the coding sequence, corroborating results obtained from ChIP-seq data.

**Fig. 4.**
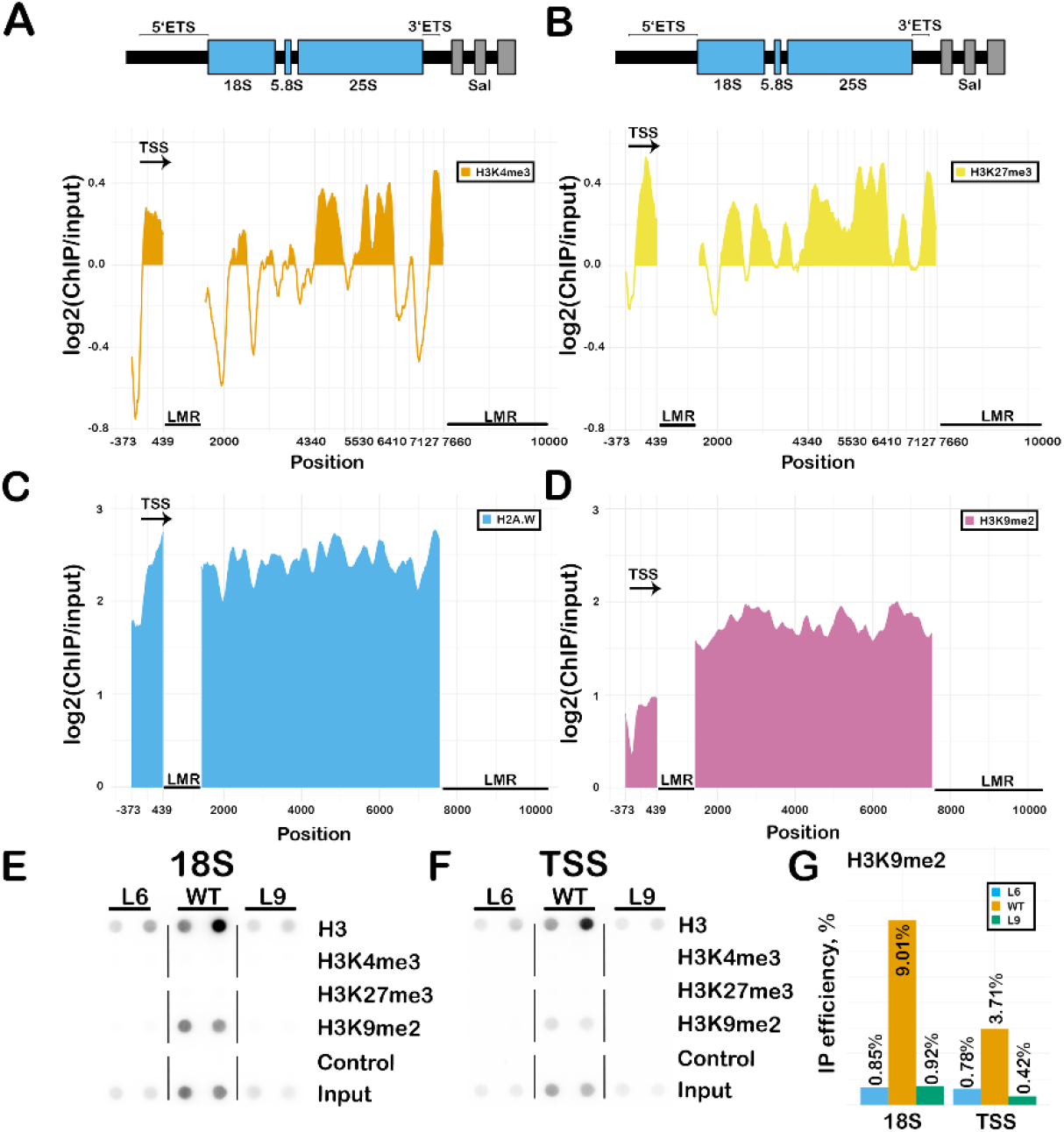
Distribution of epigenetic modifications in the rDNA. (A and B) Schematic depiction of a rDNA unit with ChIP-seq profiles of (A) H3K4me3, (B) H3K27me3, (C) H2A.W and (D) H3K9me2 in rDNA created from publicly available datasets. ChIP experiments with subsequent dot-blot signal detection showing levels of H3 histone and corresponding epigenetic marks at the (E) 18S rDNA and (F) TSS regions of a rDNA unit in WT and low rDNA copy lines L6 and L9. (G) Bar plot evaluation of H3K9me2 immunoprecipitation efficiency calculated from E and F experiments. ETS – external transcribed spacer; LMR – low mappability region; TSS – transcription start site.

### Epigenetic profile of the rDNA cluster

We next analysed whether NF accumulate any specific histone modifications using immunofluorescence (IF) detection. We found that large NF were predominantly marked by H3K9me2 (Fig. 5 *A1* and *A2*), similarly to the nuclear chromocenters while some of the smaller NF did not show H3K9me2 accumulation (Fig. 5 *A3* and *A4*). In WT, H3K9me2 labelling always occurred in perinucleolar chromocenters and in approximately 50% of NF (22 / 44; Fig. 5*B*). Low-copy line L6 showed smaller proportion of NF labelled by H3K9me2 (8 / 32; Fig. 5*B*). We further investigated whether the H3K27me3 and H3K4me3 also occur in the nucleoli. Regardless of a large number of studies containing some H3K27me3 and H3K4me3 IF data on plant nuclei (Lawrence et al, 2004; Mathieu et al, 2005), the presence of these two histone marks has neither been clearly shown nor excluded. In isolated nuclei, both H3K4me3 and H3K27me3 marks display typical diffuse distribution in the nucleoplasm (*SI appendix*, Fig. S7 *A* and *B*). We reasoned that even if these marks contribute to rDNA transcriptional regulation, they represent a relatively small proportion of total rDNA repeats. With dSTORM microscopy we found that some nucleoli appear mostly devoid of these modifications while others show modest localisation of H3K4me3 and H3K27me3 inside nucleoli (Fig. 5 *C-F*). In this experiment, we used HILO illumination and stringent settings for localisation detection, excluding localisations with low localisation precision or low emission strength to minimize the detection of the signals coming from out-of-focus fluorophore blinking.

**Fig. 5.**
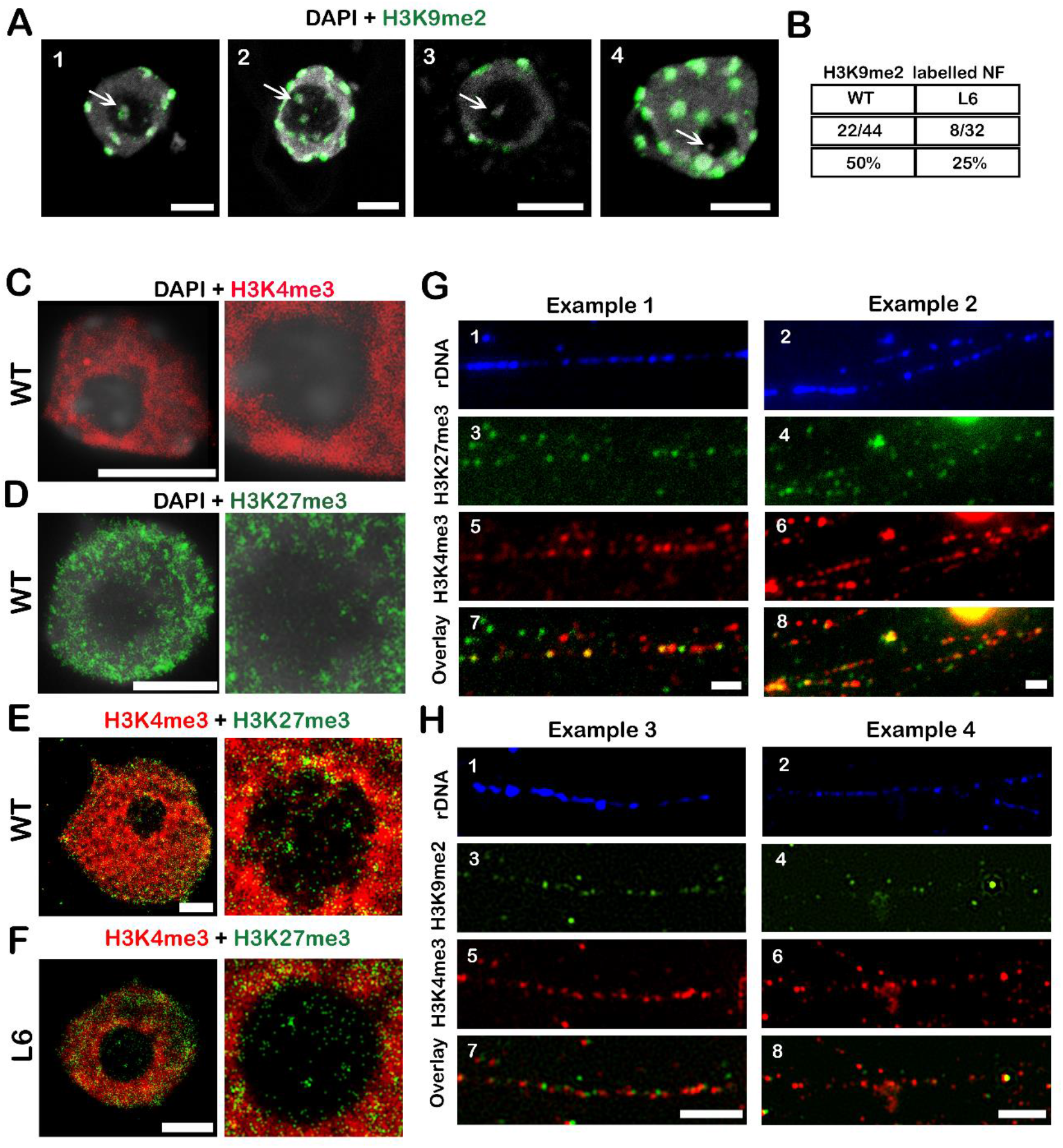
Distribution patterns of H3K9me2; H3K27me3 and H3K4m3 in rDNA based on advanced microscopy methods (A,G – confocal, C,D,E,F – dSTORM, H – SIM) (A) H3K9me2 pattern in nuclear chromocenters and nucleolar foci (NF). NF (indicated by arrows) showing either accumulation of H3K9me2 (1,2) or lack of H3K9me2 signal (3,4). (B) Percentage of nuclei with NF labelled by H3K9me2 in WT and low-copy line L9. (C-F) Images of nuclei obtained by dSTORM microscopy, insets show a more detailed view of the nucleolus. Distribution of H3K4me3 (C) and H3K27me3 (D) in plant nuclei. (E and F) Mutual distribution of H3K4me3 and H3K27me3 in the nucleoli of WT and low-copy plant lines. (G and H) Distribution of H3K4me3 (G5 and G6; H5 and H6), H3K27me3 (G3 and G4) and H3K9me2 (H3 and H4) in clusters of rDNA (G1 and G2; H1 and H2) visualized on chromatin fibers. (G7 and G8) Overlay images showing partial colocalisation between H3K4me3 and H3K27me3. (H7) Lack of colocalisation between H3K4me3 and H3K9me2 histone marks. (H8) Segments of rDNA clusters largely devoid of repressive marks. Scale bars (A) −5 μm. Scale bars (C-H) – 2 μm. NF – nucleolar foci.

In order to explore mutual positions of histone modifications along the rDNA repeat, we optimised the method of extended chromatin fibers and investigated selected histone modifications in WT plants. First, we validated FISH detection of rDNA in chromatin fiber preparations (*SI appendix*, Fig. S7*C*). Using combined FISH/IF on chromatin fibers, we detected rDNA clusters entirely devoid of the studied histone marks (*SI appendix*, Fig. S8) or observed rDNA clusters with varying arrangement of repressive H3K27me3 and active H3K4me3 marks (Fig. 5*G*). As these marks sometimes colocalized along one fiber they are likely to correspond to the bivalent chromatin regions. Repressive mark H3K9me2, in contrast, did not colocalize with H3K4me3, as expected (Fig. 5*H*). Surprisingly, we did not notice clear boundaries separating the repressive and active chromatin states (Fig. 5*H(7)*).

## Discussion

### Perinucleolar chromocenters and nucleolar foci

Chromocenters are nuclear domains where silent, condensed and heavily methylated heterochromatic parts of the genome accumulate (Fransz et al, 2006; Soppe et al, 2002). Nearly 50% of ribosomal genes are typically silenced and reside in these chromocenters (Fransz et al, 2002; Fransz et al, 2006; Pavlistova et al, 2016; Pontvianne et al, 2013; Soppe et al, 2002; Tessadori et al, 2007). In the Col-0 *Arabidopsis* ecotype, inactive NOR2 assembles into chromocenters at the nuclear periphery (Dvorackova et al, 2018; Pavlistova et al, 2016) while the active NOR4 forms chromocenters at the periphery of the nucleolus (Fransz et al, 2002; Pontvianne et al, 2013). Active rDNA genes loop into the nucleolus from perinucleolar chromocenters in a decondensed form (Highett et al, 1993; Leitch et al, 1992), reminiscent of euchromatin loops observed in the nucleoplasm (Fransz et al, 2002). Therefore, the presence of both H3.1 and H3.3 histone variants in the perinucleolar rDNA-containing chromocenters possibly reflects the complexity of rDNA organisation at NOR4, where both active and inactive copies coexist. We focused our attention on the fact that chromocenter-like structures, here called nucleolar foci (NF), occur inside the nucleolus. These foci differ in size and chromatin structure and correspond to rDNA (Fig. 1*D* and 2*D*). Although the presence of NF is evident from many cytological studies and the idea of active nucleolar rDNA being interconnected with condensed sites was previously introduced (Highett et al, 1993; Neves et al, 2005), it has not been much explored. We show that NF occur more often in the nuclei with a higher ploidy level, while nuclei of the dividing zone show nucleolar rDNA signals lacking apparent condensed sites (Fig. 2C). Interestingly, we found that the epigenetic profile of all NF is not identical. Larger foci are brightly stained with DAPI and either contain both H3 variants and H3K9me2 or they lack H3K9me2 (Fig. 5*A*). In some cells, we identified a subpopulation of NF that seemed to contain only H3.3. The co-occurrence of H3.1 histone variant and H3K9me2 is indicative of silent chromatin while H3.3 was shown to coexist with euchromatin marks, such as H3K4me3, H3K36me3 and histone acetylation (Roudier et al, 2011; Sequeira-Mendes et al, 2014). In agreement with the general idea of H3K9me2 distribution inside the inactive rDNA, summarised in (Neves et al, 2005), fewer H3K9me2 labelled NF occurred in lines with reduced rDNA copy number, where the overall abundance of H3K9me2 was also lower (Fig. 4 *E* and *F*; 5 *A* and *B*). This supports the model that inactive rDNA copies assemble into a heterochromatic state not only at the perinucleolar chromocenters but also inside the nucleolus (at NF) and that their epigenetic profile is similar. Nevertheless, NF lacking the H3.1 variant and H3K9me2, represent another type of chromatin structure. When viewed in isolation, the occurrence of H3.3 in NF could be interpreted as a sign of active transcription, facilitating the access of RNA pol I to the promoter and allowing flexible epigenetic regulation. We suppose that H3.3 presence in rDNA chromocenters is necessary for its rapid regulation, akin to environmental response genes, as shown previously (Liu et al, 2014; Wollmann et al, 2017).

Whether the condensed H3.1 and H3.3 foci inside the nucleolus truly label transcriptionally active rDNA or relate to the folding of rDNA copies is unclear. In differentiated cells, occurrence of H3.3 labelled NF correlated with rDNA transcription, as concluded from actinomycin-D (ActD) treatment leading to the depletion of H3.3 from the nucleolus (Shi et al, 2011). However, ActD was shown to induce nucleolar stress response (van Sluis & McStay, 2019), during which intranucleolar rDNA migrates to the periphery of the nucleolus (Franek et al, 2016; Harding et al, 2015). Therefore, the removal of H3.3 from nucleoli after ActD does not necessarily relate to the inhibition of RNA pol I. We analysed the root meristem, where most cells are diploid and the rDNA copy number is not amplified by polyploidization. H3.3 chromocenters were present in the plants containing WT rDNA copy number and absent in low rDNA copy lines, where the majority of rDNA copies is active (Fig. 2*A*). Meristem nuclei of low rDNA copy lines contained only the nucleolar and decondensed rDNA fraction suggesting that the formation of H3.3 chromocenters might be connected to the folding of non-transcribed rDNA and possibly correspond to the semi-condensed rDNA state, as suggested previously (Highett et al, 1993). Folding of rDNA as a mechanism of rDNA dosage control is a generally accepted concept, providing another layer of epigenetic regulation (Highett et al, 1993; Chen et al, 1998). The complexity of nucleolar rDNA, however, often impedes the visualization of the active rDNA due to the signals from condensed repeats. STORM, as we show, could bring a new insight into the rDNA folding process as it enabled detection of highly decondensed genes even in the nuclei where the nucleolus is small and rDNA condensed sites dominate (*SI appendix*, Fig. S9).

The super-resolution STED microscopy is another suitable and less demanding approach to capture the heterogeneity of the types of rDNA organization and different degrees of rDNA condensation. It revealed the presence of smaller rDNA signals dispersed in the nucleolus (Fig. 2*D*) together with NF. We suggest that the ratio of active to inactive ribosomal copies is the primary determinant of the rDNA organization inside of nucleoli. Once the requirements for rDNA transcription decrease, the folding of unnecessary repeats into NF or perinucleolar rDNA chromocenters is initiated. This explains why both types of structures occur also in the low-copy line, particularly in the endocycling root cells (*SI appendix,* Fig. S2 *D*). Terminal stage of the “packing” process is represented by the formation of a few large perinucleolar chromocenters with or without NF (Fig. 2 *D*7 and *D9*). The ratio of the nucleolar volume to the remaining nucleoplasm itself represents an indicator of the transcriptional activity of the nucleolus (Derenzini et al, 1988a; Watanabe-Susaki et al, 2014). We observed a connection between the size of nucleoli and rDNA distribution. This mostly manifests in large nucleoli, where small dispersed rDNA foci are common, and small nucleoli, where large rDNA foci are formed on the periphery of the nucleolus. Nevertheless, the regulation of rDNA folding remains elusive. Why rDNA sometimes assembles into NF while at other times it forms perinucleolar chromocenters is an open question and it is currently unknown what molecular players are mediating this process.

We reasoned that several chromatin subtypes might exist in different NF. Besides H3.3, we did not identify any active epigenetic marks in NF that would support the existence of chromocenters with uniquely active chromatin characteristics. The H3K4me3 signal was low and dispersed, with no signs of localised foci. H3K27me3, that could potentially indicate the flexible regulation of a subset of nucleolar rDNA, was also low and not associated with chromocenters (Fig. 5).

### The role of histone modifications in rDNA organization and transcription

In *Arabidopsis,* nine chromatin states, defined by different levels of DNA methylation, histone post-translational modifications and histone variants were described by genome wide analyses (Roudier et al, 2011; Sequeira-Mendes et al, 2014). Given the complexity of rDNA organisation, identification of individual chromatin states inside its large repetitive clusters is challenging. The same way the clusters of inactive rDNA copies on the nucleolar periphery obscure the microscopic analysis of the active rDNA fraction, active and poised rDNA fractions are masked by inactive rDNA in ChIP-seq analysis. IF data and ChIP-PCR show that two repressive marks H3K9me2, H3K27me1 are the most abundant modifications in nuclear chromocenters, including perinucleolar rDNA sites (Earley et al, 2010; Jacob & Michaels, 2009). The ChIP-seq analysis revealed that besides H3K9me2, H2A.W is also strongly enriched all over the rDNA repeat (Fig. 3 *C* and *D*). The broad distribution pattern of H2A.W and H3K9me2 seems characteristic for plants as in human and mice rDNA, repressive modifications form separate peaks (Zentner & Henikoff, 2014). *Arabidopsis* H2A.W is known to promote higher order chromatin condensation and it can be deposited independently of H3K9me2 (Yelagandula et al, 2014). Therefore, there might be several epigenetic mechanisms regulating the activity of rDNA. The exact nature of the molecular switch between active and silenced states in rDNA is unknown, but it does involve active and repressive histone marks (Lawrence et al, 2004; Pontes et al, 2007; Probst et al, 2004). Activation of silenced rDNA copies is known to be mediated by histone acetylation and DNA demethylation (Earley et al, 2010; Chen & Pikaard, 1997). One layer of specific rDNA regulation is ensured by the nucleolar localisation of histone modifiers, including histone deacetylase HDA6 or the histone methyltransferase SuvR4 (Thorstensen et al, 2006) (Pontvianne et al, 2013). The latter introduces H3K9me2 and its presence in the nucleolus could relate to the occurrence of H3K9me2 labelled NF (Fig. 5 *A*). Why some histone modifiers and histone marks are found in the nucleolus while others are not, remains unclear and the complete view of rDNA regulation is still missing. For example, nucleolar localisation was shown for H3S10 phosphorylation in tobacco cells (Granot et al, 2009), though it is known to associate with the mitotic progression (Demidov et al, 2009). On the contrary, two euchromatic marks H3K4me3 and H3K27me3 (active and repressive, respectively) were not clearly visible in the nucleolus (Mathieu et al, 2003; Wollmann et al, 2017). In dSTORM experiments, we could detect intranucleolar localisation of these marks (Fig. 5 *C*-*F*) and low signal emerged also in the ChIP–seq data (Fig. 4 *A* and *B*). IF-FISH on chromatin fibers, in comparison, showed quite homogeneous distribution of H3K4me3 along the rDNA clusters (Fig. 5 *G* and *H*). The use of extended chromatin fibers, as we show, helps explore the potentially hidden rDNA epigenetic patterns and provides high resolution image of the mutual distribution of individual histone modifications. We also detected H3K9me2 foci well separated from H3K4me3 sites and colocalisation between H3K4me3 and H3K27me3 (Fig. 5 *G* and *H*). The initial model of chromatin organisation in *Arabidopsis* considered H3K4me3 and H3K27me3 as mutually exclusive (Roudier et al, 2011; Schwartz & Pirrotta, 2007), but more recent studies revealed that these two marks can coexist and occur in stress response genes (Liu et al, 2014; Sequeira-Mendes et al, 2014). This corresponds with a view that bivalent (poised) chromatin allows for a more dynamic transcriptional regulation, both in the case of differentiation (Bernstein et al, 2006) and environmental response (60). It is therefore evident that several different epigenetic states exist in rDNA, shown here and in (Earley et al, 2010; Pontvianne et al, 2010; Pontvianne et al, 2013; Probst et al, 2004)

Apart from the regulation mediated via post-translational modification, histone variants contribute to the regulation of rDNA transcription. In RNA pol II transcribed genes, H3.3 is known to associate with active genes, affect gene body methylation and to be enriched in transcription end site (Ahmad & Henikoff, 2002; Stroud et al, 2012; Wollmann et al, 2017). H3.3 distribution pattern in rDNA differs from RNA pol II transcribed sites: it shows H3.3 enrichment in the TSS and lack of increase towards the 3’ end (Fig. 3 *C* and *D)*. We suggest that H3.3 in the TSS could be characteristic for RNA pol I transcribed genes and that rDNA promoter is an important regulatory element. In addition to H3.3, both H3K4me3 and H3K27me3 displayed a peak at the TSS while the levels of H3K9me2 dropped in this region. Another Pol l feature was a strong depletion of all studied histone marks before the TSS. This is reminiscent of the mouse and human rDNA, regulated via upstream control element (UCE), where a narrow peak of activating modifications was reported (Zentner & Henikoff, 2014; Zentner et al, 2011).

Assigning rDNA to any known chromatin state is not straightforward. A large subset of rDNA repeats seems reminiscent of chromatin state 9, characterised by high levels of H3K9me2, H3K27me1, CG methylation and H3.1 histone variant (Sequeira-Mendes et al, 2014). Smaller subset of rDNA is similar to the chromatin state 2, due to the co-occurrence of H3K4me3 and H3K27me3 (Sequeira-Mendes et al, 2014). The distribution of H3K4me3 and H3K27me3 fits into the model where chromatin states form only short, several kb long segments interspersed with each other (Roudier et al, 2011) and it might represent a poised rDNA state. State 2 is typical for promoters and intergenic regions (Sequeira-Mendes et al, 2014), but whether this is the case for rDNA is currently unclear. Moreover, our data indicate features that make rDNA chromatin signature unique, such as H3.3 abundance in rDNA perinuclear chromocenters and nuclear foci or H3.3 specific peak at the TSS. Interestingly, we observe presence of H3K9me2 in nucleolar foci that are universally labelled with H3.3, suggesting the existence of another chromatin environment. It is therefore evident that rDNA exists in a variety of chromatin states.

Overall, this paper uses new methodical approaches to tackle long-stranding questions in the organization and epigenetic regulation of ribosomal DNA. We show that both H3.1 and H3.3 histone variants occupy the majority of nucleolar foci and perinucleolar chromocenters, some of these sites are clearly repressive in nature since they display H3K9me2 accumulation. Importantly, we employed low-copy rDNA lines to demonstrate that nucleolar foci are formed by the compaction of rDNA in nucleoli, a process initiated once the ratio of active and inactive copies reaches a certain threshold. We captured the heterogeneity of rDNA arrangements and folding in various cell types using super-resolution microscopy and show that intermediate levels of rDNA compaction occur, besides the often-referenced clustering of rDNA signals on the nucleolar periphery. Finally, we corroborate data from previous research showing the role of active (H3K4me3) and repressive (H3K27me3; H3K9me2; H2A.W) histone marks and variants in the control of rDNA activity. We suggest that integrating various methodical avenues in the study of rDNA physiology is required given the technical hurdles of individual methods and the complexity of rDNA regulatory processes which tie together spatial organization with changes in histone signature and transcriptional activity. In conclusion, this paper presents a complex investigation of rDNA physiology, from its organization in different cell types to the epigenetic variability of rDNA clusters. Key remaining open questions are a) which histone writers and histone readers are directly responsible for toggling on and off active and inactive states in rDNA b) why is rDNA preferentially folded into perinucleolar chromocenters in some cases and into nucleolar foci in others c) why is H3.3 histone recruited into rDNA condensed foci and whether its accumulation is the result of transcription or related to a different physiological process.

## Methods

### Isolation of nuclei and nucleoli; chromatin fiber preparation

Isolation of nuclei was performed as described in (Jackson et al, 1998). Isolated nuclei were centrifuged at 5000 *g* for 10 min at 4°C and resuspended in 75 mM KCl. Chromatin fibers were extended as described in (Cohen et al, 2009), with minor modifications. Full protocol can be found in *SI appendix, Methods*. Nucleoli from 10-d-old seedlings were isolated based on the published protocol (Liang et al, 2012) with some modifications. Seedlings were fixed in 1% paraformaldehyde (PFA) for 10 min, quenched with 0.125 M glycine and disrupted in Galbraight’s buffer using ICA T25 Ultra Turrax homogenizer (11000 rpm, 90 s). After filtration through 40 μm cell strainers (Falcon, #352340), nuclei were sonicated (Diagenode Bioruptor; 5 × 5 min cycles of 30 s ON / 30 s OFF) and nucleoli were isolated by centrifugation through sucrose cushions as described.

### Immunofuorescence (IF) and FISH on nuclei and nucleoli, whole-mount FISH

Immunofluorescence detection and FISH procedures on isolated nuclei and nucleoli is described in *SI appendix, Methods*. For whole-mount FISH,10-d-old seedlings were fixed for 30 min in 1% formaldehyde, 10% DMSO, 0.5 mM EGTA in 1x PBS. Seedlings were rinsed twice in ethanol and 100% methanol, then left in methanol O/N. Seedlings were washed 2 × 5 min in 1x PBS/0.1% Tween-20 and 3 × 5 min in 2x SSC. Seedlings were incubated 30 min in the mix of 2x SSC and HB50, 1:1 (HB= hybridization buffer, 10% dextran sulfate, 50% deionized formamide in 2x SSC); 30 min in HB50 and 1-2 hours (37°C, 300 rpm) in 100 μL of the probe mix (5 μL of labelled rDNA probe in HB50). Seedlings were denatured for 4 min at 90°C, cooled on ice for 3 min and hybridized O/N at 37°C. All subsequent washes were performed in the thermomixer (300 rpm): 2 × 5 min in 50% formamide / 2x SSC at 42°C; 1 × 5 min in 2x SSC and 1 × 5 min in 1x PBS, both at 42°C. Seedlings were then put onto a microscopic slide and covered with DAPI, 2μg/mL in Vectashield.

### Chromatin immunoprecipitation (IP) and high throughput sequencing

Nuclei isolation was performed using 2 g of 10-d-old seedlings. Nuclei were sonicated, 4 × 5 min cycle (30 s on/ 30 s off; Diagenode Bioruptor) with 4 min incubation on ice between sonication cycles. Samples were then centrifuged 4 × at 20000 *g* for 10 min at 4° C. Chromatin was pre-cleared by O/N incubation with 40 μL of Protein A/G beads (sc-2003, Santa Cruz biotechnology) on a stirring wheel at 4°C. We prepared bead-antibody complexes by 4 h incubation of protein A/G agarose beads with 2 μg of antibody (anti-histone H3 ab1791; anti-histone H3K4me3 ab8580; both Abcam) in total volume of 500 μL of dilution buffer followed by O/N blocking of antibody-bead complexes. DNA obtained after the protocol was processed by QIAquick PCR purification kit (Qiagen).

### Dot-blot, hybridization and nick-translation labelling of rDNA

Dot-blot was performed according to previously described protocol (Mozgova et al, 2010). Primers and data evaluation are described in *SI appendix, Methods*. Probe labelling was adapted from (Mandakova & Lysak, 2008) using 1 μg of BAC as a template, isolated using Macherey Nagel NucleoBond® Xtra Midi. For visualization of the 45S rDNA loci the BAC clone T15P10 (GenBank AL095897/8) was used.

### FLIM-FRET

10-d-old seedlings were transferred from growth plates onto microscopic slides and imaged using Zeiss LSM 780 (Carl Zeiss MicroImaging GmbH) microscope equipped with “In *Tune*” laser (488 nm for GFP, 561 nm for mRFP). FLIM was performed at 488 nm and 40 MHz frequency and recorded using HPM 100-40 (Becker&Hickl, Berlin, Germany) hybrid detector, SPC-150 TCSPC (Becker&Hickl) module and SPCM64 software. Lifetime images were further processed and analyzed in SPCImage 6.4 software (Becker&Hickl).

### Microscopy and image analysis

Whole-mount FISH and chromatin fiber images were acquired on an epifluorescence microscope Zeiss AxioImager Z2, using appropriate filters. Structured illumination microscopy was performed on Nikon 3D N-SIM microscope (inverted Nikon Eclipse Ti-E, Nikon) equipped with a Nikon CFI SR Apo TIRF objective (100x oil, NA 1.49). Two-color STED microscopy was performed on the Abberior STED system joined to a Nikon Eclipse Ti-E inverted confocal microscope. For dSTORM, samples labelled with either AF647 or CF680 (1:100 secondary antibody dilution) were imaged in the imaging buffer (50 mM Tris-HCl, 10 mM NaCl, 10% (w/v) glucose, pH 8 with 50 mM β-mercaptoethylamine (MEA, 30070, Sigma-Aldrich), 1.1 mg/ml Glucose Oxidase (G2133, Sigma-Aldrich) and 100 μg/ml Catalase (C40, Sigma-Aldrich).

Detailed description of plant material, growth conditions and all methods is presented in *SI appendix, Methods*.

## Acknowledgements

We are grateful to: C. Gutierrez and S. Otero CBMSO, Madrid, Spain, for providing HTR5-mRFP/HTR3-GFP, HTR4-myc and HTR3-myc plant lines; B. Desvoyes CBMSO, Madrid, Spain, for critical reading of the manuscript and fruitful discussions; F. Pontvianne, CNRS, Perpignan, France, for providing the FIB2-YFP plant line; A. Probst, GReD, Clermont-Ferrand for the help with ChIP experiment optimization; D. Pánek, Imaging Methods Core Facility, for the support with the realization of spectral de-mixing experiment and analysis. We would like to acknowledge these core facilities (CF) for their support with obtaining data presented in this paper: CELLIM of CEITEC MU and Imaging Methods CF at BIOCEV, both supported by MEYS CR (LM2018129 Czech-BioImaging); Plants Sciences and Genomics of CEITEC MU.

MD and MF were supported by the Czech Science Foundation project 19-11880Y, by Ministry of Education, Youth and Sports of the Czech Republic - project INTER-COST (LTC18048) and by the European Regional Development Fund - Project “SINGING PLANT” (CZ.02.1.01/0.0/0.0/16_026/0008446). MO acknowledges financial support from the ERDF (project No. CZ.02.1.01/0.0/0.0/16_013/0001775).

## Author Contributions

KK, MF and MD performed the experiments and wrote the manuscript. MO assisted with super-resolution microscopy and subsequent image analysis. KD and JK assisted with ChIP-seq data analysis.

## Supplementary information

### Supplementary material

#### Plant material

All *A. thaliana* lines used in this study were derived from the Columbia ecotype. Previously described lines were used: *fas1-4* (At1g65470) (NASC: N828822, SAIL_662_D10 (Exner et al, 2006; Mozgova et al, 2010); plant lines transformed with HTR3-GFP, HTR4-GFP, HTR3-Myc, HTR4-Myc, HTR13-mRFP and double transformants HTR5-mRFP/HTR3-GFP (“RedGreen”) were provided by C. Gutierrez CBMSO, Madrid, Spain (Otero et al, 2016; Stroud et al, 2012); fibrillarin-YFP plant line was obtained from F. Pontvianne, CNRS, Plant Genome and Development, Perpignan, France. Plant lines with reduced rDNA amount – line 6 (L6) and 9 (L9) were previously obtained and characterised in our laboratory (Pavlistova et al, 2016).

In *A.thaliana*, H3 variants are encoded by several HISTONE 3 RELATED (HTR) genes, namely HTR1, HTR2, HTR3, HTR9 and HTR13 for H3.1 and HTR4, HTR5 and HTR8 encoding H3.3.We used HTR4 (At4g40030) and HTR5 (At4g40040) for H3.3 and HTR3 (At3g27360) and HTR13 (At5g10390) for H3.1. Plant line carrying both HTR4-GFP and HTR13-mRFP (“GreenRed”) fusion proteins was obtained by crossing of HTR4-GFP and HTR13-mRFP plants and selection of double labelled plants in F2. HTR4-GFP + L9 were prepared by crossing the *fas1-4* expressing HTR4-GFP (Otero et al, 2016) into line L9 background (Pavlistova et al, 2016).

### Supplementary methods

#### Plant growth

All seeds were sterilized (70% ethanol/ 10 min followed by 99% ethanol/ 5 min) and plated on half-strength agar Murashige and Skoog medium (½ MS medium) with 1% sucrose. After 2-day stratification (4°C/dark) plates were transferred to the growth chamber and pre-grown up to 2 weeks under long day (LD) conditions (16 h light −21°C/8 h dark −19°C/50-60% relative humidity). Seedlings were then collected directly from plates and used for experiments or transferred into the soil and grown under the same long day conditions or transferred to the short day (8 h light −21°C/16 h dark −19°C/50-60% relative humidity).

#### Isolation of unfixed nuclei for fiber preparations

Twelve-d-old seedlings (0.5 g) were chopped with the razor blade in ice-cold nucleus isolation buffer (NIB −0.5 M sucrose; 10 mM EDTA; 2.5 mM DTT; 100 mM KCl; 1 mM spermine; 4 mM spermidine in 10 mM Tris‐Cl, pH 9.5). After chopping, the solution was filtered through 50 μm and 30 μm pore size filters (CellTrics, Sysmex, Germany). Afterwards, the filtrate was supplemented with 1/10 volume of 10% Triton‐X in NIB and centrifuged at 2000 *g* for 10 min at 4°C. The pelleted nuclei were then resuspended in 400 μL of NIB and supplemented with 400 μL of 100% glycerol. Nuclei were then aliquoted and stored at −20°C.

#### Isolation of nucleoli

Nucleoli from 10-d-old seedlings were isolated based on the published protocol (Liang et al, 2012) with some modifications. Seedlings were fixed in 1% paraformaldehyde (PFA) for 10 min then quenched with 0.125 M glycine for 5 min (both under vacuum). Seedlings were disrupted in 20 mL of Galbraith buffer, (GB), without Triton × (Galbraith et al, 2011) using ICA T25 Ultra Turrax homogenizer (11000 rpm, 90 s) and filtered through 40 μm nylon cell strainer filters (Falcon, #352340). Then Triton-X was added to the final concentration of 0.3%. Homogenate was vortexed, left 5 min on ice and centrifuged at 350 *g* for 20 min at 4°C. Pellet was resuspended in Solution I (Liang et al, 2012) (0.5 M sucrose with 3 mM MgCl2 and protease inhibitors), 500 μL aliquots were used for sonication (Diagenode Bioruptor; 5 × 5 min cycles of 30 s ON / 30 s OFF at H mode). Sonicated nuclei were underlaid with 700 μL of Solution II (1 M sucrose, 3 mM MgCl2 with protease inhibitors) and centrifuged at 1800 *g* for 5 min at 4°C. Pellet containing nucleoli was resuspended in 500 μL of storage buffer (90% Glycerol, 10% GB) and stored at −20°C.

#### Chromatin fiber preparation

An aliquot of isolated nuclei was centrifuged at 5000 *g* for 10 min at 4°C. Then, nuclei were resuspended in 75 mM KCl (the dilution needs to be optimized for each isolation, for chromatin fibers we recommend first testing 1:50; 1:100 and 1:200 dilutions) and incubated for 20 min. 8 μL of the nuclear suspension were cytocentrifuged onto poly-L-lysine-coated coverslips at 400 *g* for 4 min at 4°C. Coverslips were immediately removed from the centrifugation chambers, excess liquid was removed and 20 μL of nucleus lysis buffer (NLB −330 mM NaCl, 500 mM urea, 1 % Triton-X 100, 25 mM Tris-Cl, pH = 7) were pipetted onto nuclei. The drop was quickly covered with a square parafilm coverslip (15×15 mm in size). NLB was left to evaporate for 60 min, after which the coverslips were immersed in 1x KCM buffer (120 mM KCl, 20 mM NaCl, 10 mM Tris-Cl, 0.5 mM EDTA, 0.1% Triton-X) for 30 min. Blocking was performed with 5% BSA in KCM for 30 min at RT, after which the samples were incubated with primary antibodies diluted 1:100 in 5% BSA / KCM solution for 1 h at RT (anti-histone H3K4me3, ab8580; anti-histone H3K27me3, ab6002; anti-histone H3K9me2, ab1220). Samples were washed three times for 5 min in KCM, then incubated with appropriate secondary antibodies diluted 1:200 in 5% BSA / KCM. Samples were washed three times for 5 min in KCM, post-fixed in 4% PFA in KCM, followed by two washes in 1x PBS. After mounting in DAPI + Vectashield, fiber stretching and immunofluorescence was checked on an epifluorescence microscope Zeiss Axioimager Z1. Antifade was then removed by two times 5 min washes in 1x PBS. Hybridisation mixture (50% deionized formamide; 10% dextran sulfate; 1:30 diluted nick-translation labelled rDNA probe) was then pipetted onto clean glass slides and the coverslips with fibers were put face-down onto the glass slides and sealed off with rubber cement. Samples were denatured at 75°C for 3 min in a thermal cycler and hybridized O/N at 37°C. The next day, coverslips were washed once in 2x SSC at RT, then 3x in 50% formamide (FA) wash buffer (50% FA, 2x SSC) at 37°C and then once in 2x SSC at 37°C. Coverslips were mounted in DAPI (4′,6-Diamidine-2′-phenylindole dihydrochloride), 2 μg/mL in Vectashield and imaged.

#### Nick-translation rDNA labelling

Probe labelling was adapted from (Mandakova & Lysak, 2008) using 1 μg of BAC as a template, isolated using Macherey Nagel NucleoBond® Xtra Midi. For visualization of the 45S rDNA loci the BAC clone T15P10 (GenBank AL095897/8) was used.

#### Immunofuorescence (IF) and FISH on nuclei and nucleoli

Half g of 10-d-old seedlings fixed in 4% PFA for 10 mins, then rinsed in GB, see above and homogenized with a razor blade in GB. The homogenate was filtered through 50 μm and 30 μm pore-size filters (CellTrics, Sysmex, Germany). Nuclei were centrifuged for 20 min at 350 *g* at 4°C, resuspended in 1x PBS and stopped onto slides. Nuclei were briefly dried at 4°C, then fixed in 4% PFA in 1x PBS / 0.5% Triton-X. Nuclei were rinsed three times in 1x PBS, blocked in 5% BSA /1x PBS for 30 min and incubated with anti-Myc antibody (M4439; Sigma Aldrich) diluted 1:100 in 5% BSA /1x PBS for 1 hour. Nuclei were washed three times for 5 min in 1x PBST (0.05% Tween-20), then incubated with an appropriate secondary antibody (1:200 dilution, 1 h incubation). Slides were washed three times for 5 min in 1x PBST, then dehydrated in ethanol series. FISH procedure was performed as described in (Dvorackova et al, 2018).

#### Whole-mount FISH

Five to ten seedlings (10-d-old) were fixed for 30 min in 5 ml of fixative (1% formaldehyde, 10% DMSO, 0.5 mM EGTA in 1x PBS). Seedlings were rinsed two times 10 min in 100% methanol and two times 10 min in 96% ethanol, transferred to a 1.5 ml Eppendorf tube and left at 4°C overnight. All subsequent washes were performed in Eppendorf tubes containing 1 ml of appropriate buffer. Seedlings were transferred to a new tube, washed two times for 5 min in 1x PBS/0.1% Tween-20 and three times for 5 min in 2x SSC. Seedlings were incubated for 30 min in the mix of 2x SSC and HB50, 1:1 (HB= hybridisation buffer, 10% dextran sulfate, 50% deionized formamide in 2x SSC); 30 min in HB50 (1 ml) and 1-2 hours (37°C, shaking at 300 rpm) in 100 μL of the probe mix (5 μL of AlexaFluor 594 labelled rDNA probe in HB50). Before the incubation in the probe mix, seedlings were briefly dried on the filter paper to avoid probe dilution and the tube was vortexed. Seedlings in the probe mix were denatured for 4 min at 90°C in a thermomixer, cooled on ice for 3 min and hybridized O/N at 37°C. All subsequent washes were performed in the thermomixer (shaking at 300 rpm): 2x 5 min in 50% formamide / 2x SSC at 42°C; 1×5 min in 2x SSC and 1×5 min in 1x PBS, both at 42°C. Seedlings were then disentangled from one another in 1x PBS in a petri dish, put onto a microscopic slide and covered with DAPI, 2μg/mL in Vectashield.

#### Chromatin immunoprecipitation (IP) and high throughput sequencing

Two grams of 10-d-old seedling were fixed in 40 ml of 1% formaldehyde and quenched with 2.7 ml of 2M glycine on ice, rinsed several times with cold mQ water, dried with paper towels and frozen in liquid nitrogen. After grinding in N2 powder was transferred in 15 ml of Extraction buffer (10 mM Tris HCl pH 8, 10 mM MgCl2, 1 M Sucrose, 5 mM β-Mercaptoethanol, 0.1 mM PMSF and protease inhibitors - PI - Pepstatin and Leupeptin at 1μg/ml final concentration) and thoroughly mixed using vortex. Solution was then filtered through 40 μm cell strainer (Falcon) and miracloth and spun down for 20 min (following centrifugation steps were performed at 1000g and 4°C). Pellet was resuspended in 5 ml of Extraction buffer supplemented with 1% Triton X-100, transferred into 15 ml tube and centrifuged for 10 min. This procedure was repeated 5 times in total. Pellet was then resuspended in 5 ml of Extraction buffer without sucrose, centrifuged for 10 min and resuspended in 300 μl of nuclei lysis buffer (50 mM Tris HCl pH 8, 10 mM EDTA, 5 mM β-Mercaptoethanol, 1% Triton X-100, 0.1 mM PMSF and PI). 10 μl of Input 1 were taken at this step for further control of genomic DNA integrity. Samples were then sonicated at diagenode bioruptor sonicator: 4×5min cycle (30 s ON/ 30 s OFF) with 4 min incubation on ice in between. Samples were then centrifuged four times at 20817 g for 10 min at 4°C and supernatant was collected into new tube each time. 20 μl of Input 2 were taken afterwards from samples. Integrity of genomic DNA and efficiency of the sonication were checked using Inputs 1 and 2 on 1% agarose gel. Sonication fragment length was optimised to 100-500 bp. Supernatant was further split into two tubes and diluted 10 times with dilution buffer (16.7 mM Tris HCl pH 8, 1.2 mM EDTA, 1.1% Triton X-100, 167 mM NaCl). Chromatin was pre-cleared using protein A/G agarose beads (40 μl per sample) that were washed with dilution buffer (300g centrifugation at 4°C) prior to incubation. Pre-clearing was performed overnight on a stirring wheel at 4°C. Bead-antibody complexes were prepared simultaneously: protein A/G agarose beads (40ul per immunoprecipitation reaction) were washed with dilution buffer and incubated with 2 μg of antibody for 4 hours on a stirring wheel. Antibody-bead complexes were then blocked overnight in blocking buffer on a stirring wheel (200 μg/ml glycogen, 3% BSA in dilution buffer). Pre-cleared samples were centrifuged at 1800g for 5 min at 4°C and supernatant was split into two tubes (equal volume). 5% of volume from each aliquote was collected together into a tube for further estimation of background (input sample). Antibody-bead complexes were washed twice with dilution buffer after blocking, mixed with samples and incubated overnight on a stirring wheel at 4°C. Washing was performed using 4 buffers: wash buffer 1 (50 mM TrisHCl pH 8, 150 mM NaCl, 1% Igepal, 0.25% Sodium deoxycholate, 1mM EDTA pH 8), wash buffer 2 (100 mM TrisHCl pH 8, 500 mM LiCl, 1% Igepal, 1% Sodium deoxycholate), wash buffer 3 (100 mM TrisHCl pH 8, 150 mM NaCl, 500 mM LiCl, 1% Igepal, 1% Sodium deoxycholate), and TE buffer according to the scheme: wash buffer 1 – wash 2 times, wash buffer 2 – 2 times, wash buffer 3 – one time, TE – 2 times. Each wash was performed with 5 min incubation on stirring wheel in a cold room followed by 5 min centrifugation at 300g at 4°C. Tubes were changed at least twice during the washes. 250 μl of elution buffer (1% SDS 0,1M NaHCO3) were added to washed beads and they were incubated at thermomixer (65 °C, 1100 rpm). Samples were centrifuged at 1800g for 5 min and supernatant was transferred into a new tube. Elution procedure was then repeated and second supernatant was mixed with the first one. 20 μl of 5M NaCl were added to supernatant followed by incubation at 65 °C overnight at 1100 rpm on thermomixer. Input samples were added at this stage of protocol. Their volume was adjusted to 500 μl using elution buffer and they were treated the same as immunoprecipitated samples. Samples were treated with RNAse by adding 20 μl of TrisHCl pH 6.5 and 5 μl of RNAse with incubation at 37 °C for 30 min, 400 rpm. This was followed by proteinase K treatment by adding 10 μl od 0.5M EDTA and 11.25 μl of proteinase K with incubation for 2 hours at 45 °C, 400 rpm. 11.25 μl of proteinase K were added afterwards with additional two-hour incubation under same conditions. Samples were precipitated by adding one tenth volume of sodium acetate 3M, pH 4.8, 1 μl of glycogen and 3 volumes of EtOH. After thorough mixing samples were left at −20 °C overnight. DNA was then pelleted by centrifugation at 20817g in a pre-cooled centrifuge for 1 hour. Pellet was then washed twice with cool 70% ethanol, each wash was followed by 15 min centrifugation at 20817g, 4 °C. Pellet was then air-dried and resuspended in 40 μl of mQ water. All samples, except the first input were then cleaned using QIAquick PCR purification kit (Quiagen) according to instructions and eluted using 40 μl of mQ water. Precipitated DNA was used for further analysis.

#### ChIP-seq data analysis

Since ribosomal genes are not well annotated in the public databases, we first created a reference genome by merging the rDNA consensus sequence (Havlova et al, 2016) with TAIR10 genome. This sequence contained coding part of rDNA, 3’ and 5’ external transcribed spacer (ETS) sequences and non-transcribed spacer (NTS) and was merged with chromosome 2 sequence. Please note that there are some rDNA repeats already in TAIR10 genome, we performed N-masking of those using bedtools maskfasta command to avoid usage of those repeats and therefore loss of signal on our reference rDNA sequence.

We then evaluated the mappability of signals coming from different parts of rDNA sequence. Mappability is a measure of the uniqueness of a given genomic sequence based on the number of its fragments that are uniquely aligned to the particular sequence (Cheung et al, 2011; Zentner & Henikoff, 2014). This analysis is necessary mostly because of NTS sequence that is not unique like the coding part and can have some homology to sequences outside of rDNA repeat. We used bias elimination algorithm for deep sequencing (BEADS) to calculate mappability values (Cheung et al, 2011). First rDNA repeat was sheared using GenomeTools shredder command into short sequences of 50 bp length that correspond to lengths of ChIPseq reads and mapped them onto the constructed genome using bowtie2 software. Mapped reads were extended to 200 bp using beads extend command, as this length corresponds to the size of DNA associated with a single nucleosome. Mappability was calculated using beads tagCount command at for each base pair of our sequence (-base 1 option). Maximum mappability score with fragment length 200 can be 400 and it is considered as 100% mappability. Signal detected at this base pair is considered to be produced exclusively by rDNA. Base pairs with mappability lower than 25% are considered poorly mappable and should not be analysed since the signal could map from some other loci in the genome. Signals of low mappability were mainly localised in NTS and the highest mappability scores were at the promoter region and at the coding sequence (*SI Appendix*, Fig.S4 *A*).

ChIP-seq reads from our experiments and public datasets were aligned using bowtie 2 to previously described genome. Further processing was done using HOMER software (Heinz et al, 2010) according to the steps described in HOMER next generation sequencing analysis tutorial. Tag directories were constructed using makeTagDirectory command with selection of uniquely alignable reads (-unique option). Data for tag distribution graphs were constructed using makeUCSCfile command and reads from experimental samples were normalized to control or input samples depending on deposited datasets. Logarithmic option –log was used for normalisation. We also selected - fragLength 200 and - inputFragLength 200 options to get data corresponding to single nucleosome.

Tag density distribution plots were visualized using R studio software (R version 3.5.2 “Eggshell Igloo”).

#### Dot-blot and hybridisation

Dot blot was performed according to previously described protocol (Mozgova et al, 2010). All immunoprecipitated DNA was alkali-transferred onto a Hybond-N+ nylon membrane (Amersham) and hybridized with α-^32^P labelled 18S rDNA probe (Mozgova et al, 2010). Signals were visualized using Fujifilm FLA7000 system. Signal intensity was analysed in Fujifilm Multi Gauge software and normalized to input signals. We used following primers for probe labelling. Probes for transcriptional start site and 18S were prepared using TSS rDNA Fwd: (AGTATCCTTATGATGCATGCCA), TSS rDNA Rev: (CCCTAACGCCTCGAAGAACTAAT

), 18S rDNA Fwd: (CTAGAGCTAATACGTGCAACAAAC) and 18S rDNA Rev:

(GAATCGAACCCTAATTCTCCG) with plant genomic DNA as a template according to (Mozgova et al, 2010).

#### Microscopy and image analysis

Whole-mount FISH, chromatin and DNA fiber images were acquired on an upright epifluorescence microscope Zeiss AxioImager Z2 using a 63x (1.40 NA) Plan Apochromat or a 100x (1.40 NA) Plan Apochromat objective, using appropriate filters. Images were captured on a Hamamatsu ORCA Flash camera with 2048×2048 pixel resolution. 3D-deconvolution on whole-mount FISH samples was conducted in ZEN software, using the slow iterative deconvolution algorithm.

#### FLIM-FRET

Fluorescence resonance energy transfer (FRET) was evaluated using fluorescence lifetime imaging (FLIM) and performed on 10-d-old *A.thaliana* seedlings using the following lines: RedGreen, GreenRed, HTR3-GFP (control) and HTR4-GFP (control). Seedlings were transferred from growth plates onto microscopic slides and imaged using Zeiss LSM 780 (Carl Zeiss MicroImaging GmbH) microscope equipped with “In *Tune*” laser (488 nm for GFP, 561 nm for mRFP). FLIM was performed at 488 nm and 40 MHz frequency and recorded using HPM 100-40 (Becker&Hickl, Berlin, Germany) hybrid detector, SPC-150 TCSPC (Becker&Hickl) module and SPCM64 software. Lifetime images were further processed and analyzed in SPCImage 6.4 software (Becker&Hickl). FRET levels were further calculated as [fluorescence lifetime(control) – fluorescence lifetime(experiment)]/fluorescence lifetime(control). Acceptor over donor ratio was calculated as [fluorescence intensity(acceptor) - fluorescence intensity (donor)]/fluorescence intensity(acceptor). Fluorescence intensity was measured with Fiji software using images of studied nuclei taken prior to FLIM measurement. ROI (region of interest) was selected (whole nucleus, nucleolar chromocenters) and mean fluorescence intensity was measured.

#### STED microscopy

Two-color STED microscopy was performed on the Abberior Instruments Expert Line STED system equipped with Nikon Eclipse Ti-E microscopy body and Nikon CFI Plan Apo Lambda 60x Oil, NA 1.40 objective. Sample was illuminated with pulsed 561 nm and 640 nm lasers and depleted by pulsed 775 nm STED laser of 2D donut shape formed by spatial light modulator. Fluorescence signal passed the pinhole set to 1AU, was filtered by emission filters (580-630 nm and 650-720 nm) and was detected with single photon counting modules (Excelitas Technologies). STED images were scanned with a pixel size of 20 nm×20 nm, 10 μs dwell time, and in line interleaved acquisition with time gated detection using the Imspector software (Abberior Instruments). DAPI was acquired with a 405 nm laser in a conventional confocal mode. For STED microscopy, rDNA was labelled with Alexa Fluor 594 and Myc-tagged histones were labelled with α-mouse Abberior Star Red secondary antibody (1:100 dilution).

Colocalisation analysis was performed in Icy image analysis software in the “Colocalisation studio” plugin(Lagache et al, 2015). For all the samples, regions of interest were selected around rDNA foci (n = 20 per condition) and Pearson R correlation coefficient and cross-correlation analysis was calculated.

#### Structured Illumination Microscopy

Structured illumination microscopy was performed on Nikon 3D N-SIM microscope (inverted Nikon Eclipse Ti-E, Nikon) equipped with a Nikon CFI SR Apo TIRF objective (100x oil, NA 1.49). Structured illumination pattern projected into the sample plane was created on a diffraction grating block (100 EX V-R 3D-SIM) for laser wavelengths 488, 561 and 647 nm. Excitation and emission light was separated by filter cubes with appropriate filter sets SIM488 (ex. 470-490, em. 500-545), SIM561 (556-566, 570-640) and SIM647 (590-650, 663-738). Emission light was projected through a 2.5x relay lens onto the chip of the EM CCD camera (Andor iXon Ultra DU897, 10 MHz at 14-bit, 512×512 pixels). Laser intensity, EM gain and camera exposure time were set independently for each excitation wavelength. Intensity of fluorescence signal was held within the linear range of the camera. 15 images (3 rotations and 5 phase shifts) were recorded for every plane and color. SIM data were processed in NIS-Elements AR. Before sample measurement, the symmetry of point spread function was checked with 100 nm red fluorescent beads (580/605, Carboxylate-Modified Microspheres, Life Technologies) mounted in Abberior Liquid Mount medium (Abberior), and optimized by adjusting objective correction collar.

#### Single Molecule Localisation Microscopy

SMLM experiments (STORM) were acquired on a N-STORM system (Nikon) equipped with Nikon Eclipse Ti body, objective Nikon CFI HP Apo TIRF 100x Oil/NA 1.49, Perfect Focus System (hardware autofocus for the stabilization of z-position) and EM CCD Andor iXon Ultra DU897 camera. For STORM, 647 nm laser beam (fiber output power 125 mW) was focused by 2x magnifying lenses in TIRF Illuminator unit to reach higher laser intensity in the sample. The sample was illuminated by highly inclined laser beam to improve the signal to noise ratio. Excitation and emission light was separated on 405/488/561/647 nm Laser Quad Band filter cube (TRF89902, Chroma), specifically, the far-red emission was collected in the emission range 674-785 nm. For spectral demixing, the microscope was adapted by insertion of the W-VIEW GEMINI Image splitting optics (A12801-01, Hamamatsu) to the right port of microscope in front of the EM CCD camera. The fluorescence emission was split to shorter and longer wavelengths by 700 nm dichroic beam splitter (FF700-Di01, Semrock). All emission wavelengths were detected by the EM CCD camera. Sequences of frames (typically 20 000-30 000 frames) were acquired in NIS-Elements software (version 5.11.02) in Fast Time-Lapse mode to capture the images with the frame rate 31.3 FPS. The time series were measured in selected region of interest with the image size of 512×128 pixels and pixel size 107 nm (the result of combination 100x/1.49 NA objective, a 1.5x zoom adapter and the camera pixel size of 16 × 16 μm^2^). Before the acquisition, the sample was illuminated by 647 nm laser to switch off most of the molecules to the dark state. The acquisition was started after detecting single molecules only. The imaging buffer (50 mM Tris-HCl, 10 mM NaCl, 10% (w/v) glucose, pH 8 with 50 mM β-mercaptoethylamine (MEA, 30070, Sigma-Aldrich), 1.1 mg/ml Glucose Oxidase from Aspergillus niger (G2133, Sigma-Aldrich) and 100 μg/ml Catalase from bovine liver (C40, Sigma-Aldrich) was freshly prepared on ice before imaging. Image reconstruction was performed in rapidSTORM software (Wolter et al, 2012) followed by drift correction based on cross-correlation analysis in ThunderSTORM (Ovesny et al, 2014). Localisations were filtered by localisation intensity and precision. For two-color images, spectral demixing of AF647 and CF680 was performed in SD-mixer software (Tadeus et al, 2015). Transformations of localisation tables between the used softwares we performed in custom writen Python scripts.

### Supplementary figures

**Fig. S1.**
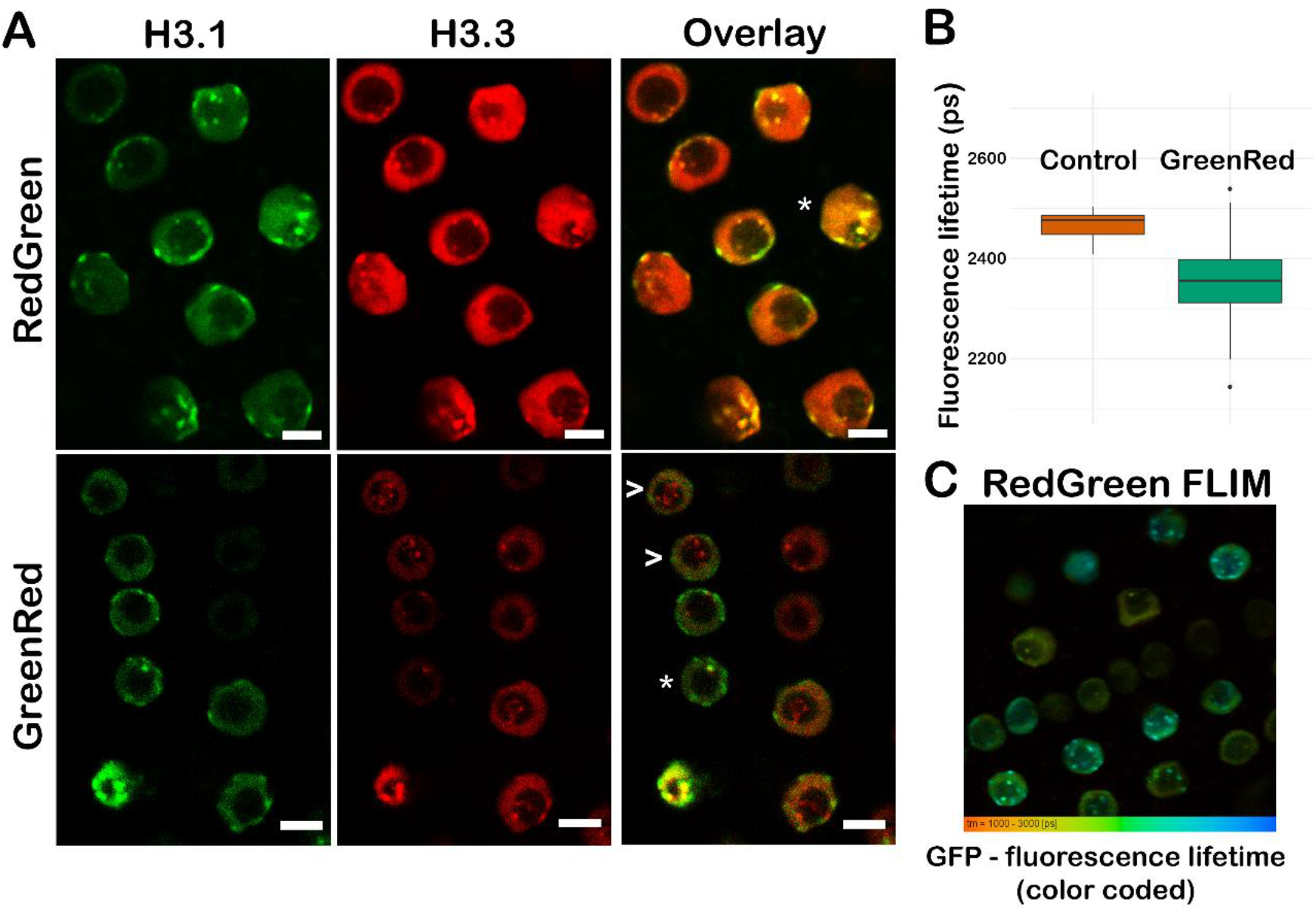
H3.1 and H3.3 distribution in root tissues. (A) Representative images of H3.1 and H3.3 distribution in root tissue for GreenRed (H3.3-GFP; H3.1-mRFP) and RedGreen (H3.3-mRFP; H3.1-GFP) plant lines. H3.1 always color-coded green and H3.3 red for clarity Nuclei with monovariant nucleolar foci are marked by an arrowhead, nuclei with bivariant nucleolar foci are marked by an asterisk. (B) FLIM-FRET experiment showing fluorescence lifetimes (τ) for control (H3.3-GFP) and GreenRed plant lines. (C) Image of root nuclei showing differences in GFP lifetime in individual nuclei. Fluorescence lifetime is color coded from red (shortest τ in red, longest τ in blue). Scale bar – 5 μm.

**Fig. S2.**
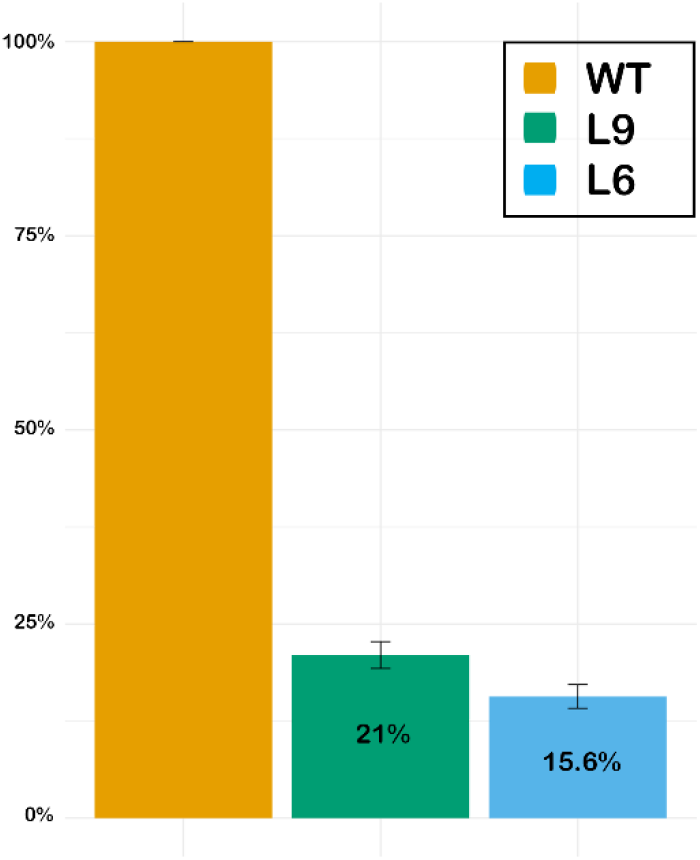
Relative 45S rDNA copy number in seedlings of WT and low rDNA copy plants (L6, L9). Relative 45S rDNA copy number of low copy lines was evaluated by qPCR in three replicas. WT sample was considered as reference for calculations.

**Fig. S3.**
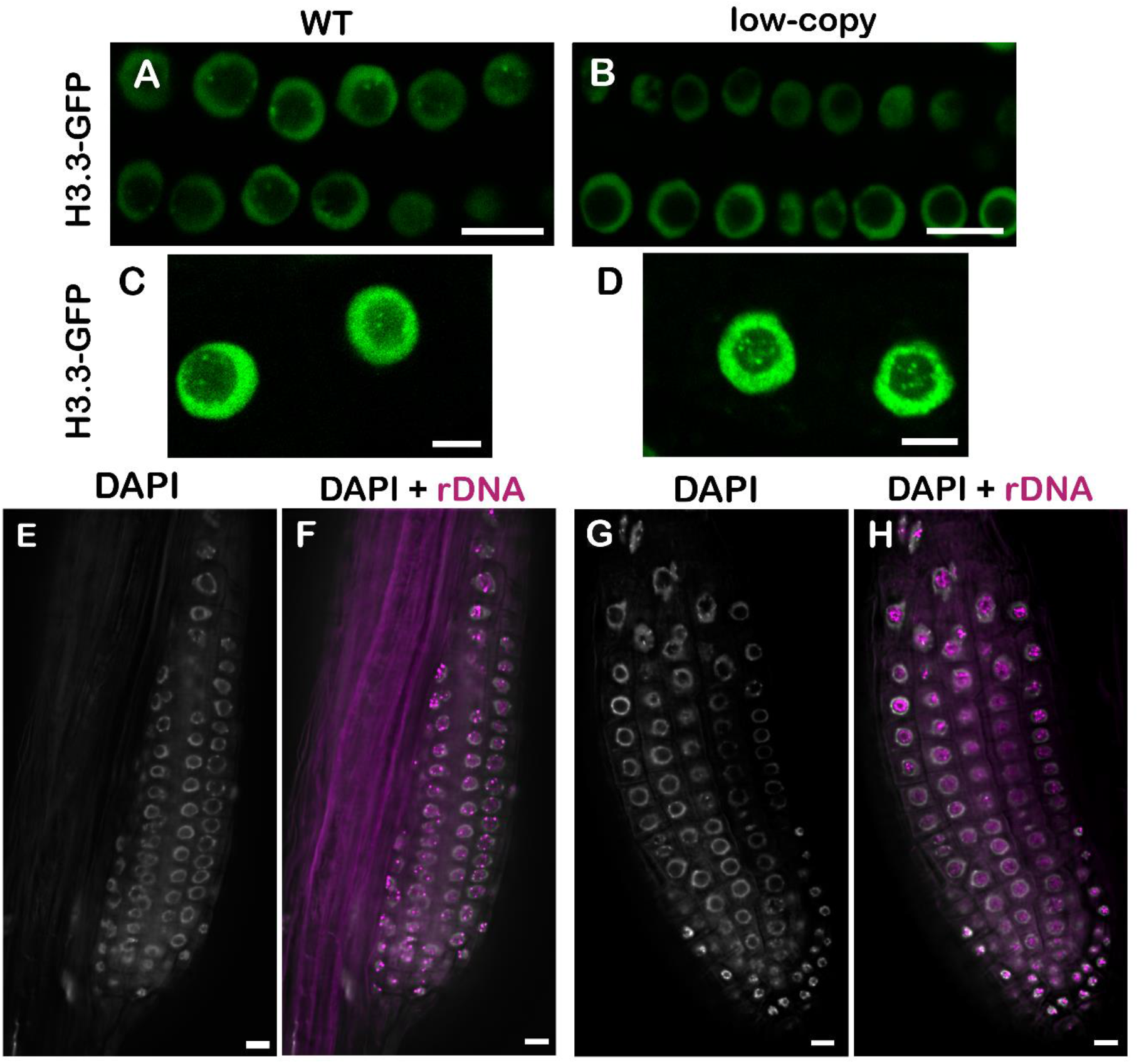
Loss of rDNA chromocenters in low-copy plants. (A and B) Distribution of H3.3-GFP in actively dividing nuclei of (A) WT and (B) low copy roots (L9 plant line), confocal images. (C and D) Examples of nuclei from the transition zone with H3.3 foci from both (C) WT and (D) low-copy line L9. (E-H) Organization of rDNA in lateral roots of (F) WT and (H) low copy plants (L6 plant line), rDNA (magenta), DAPI (grey, images E and G), wide-field microscopy combined with deconvolution. Note the absence of perinucleolar rDNA foci in the low-copy plant line (H). Scale bar – 5 μm (A and B); 10 μm (C-H).

**Fig. S4.**
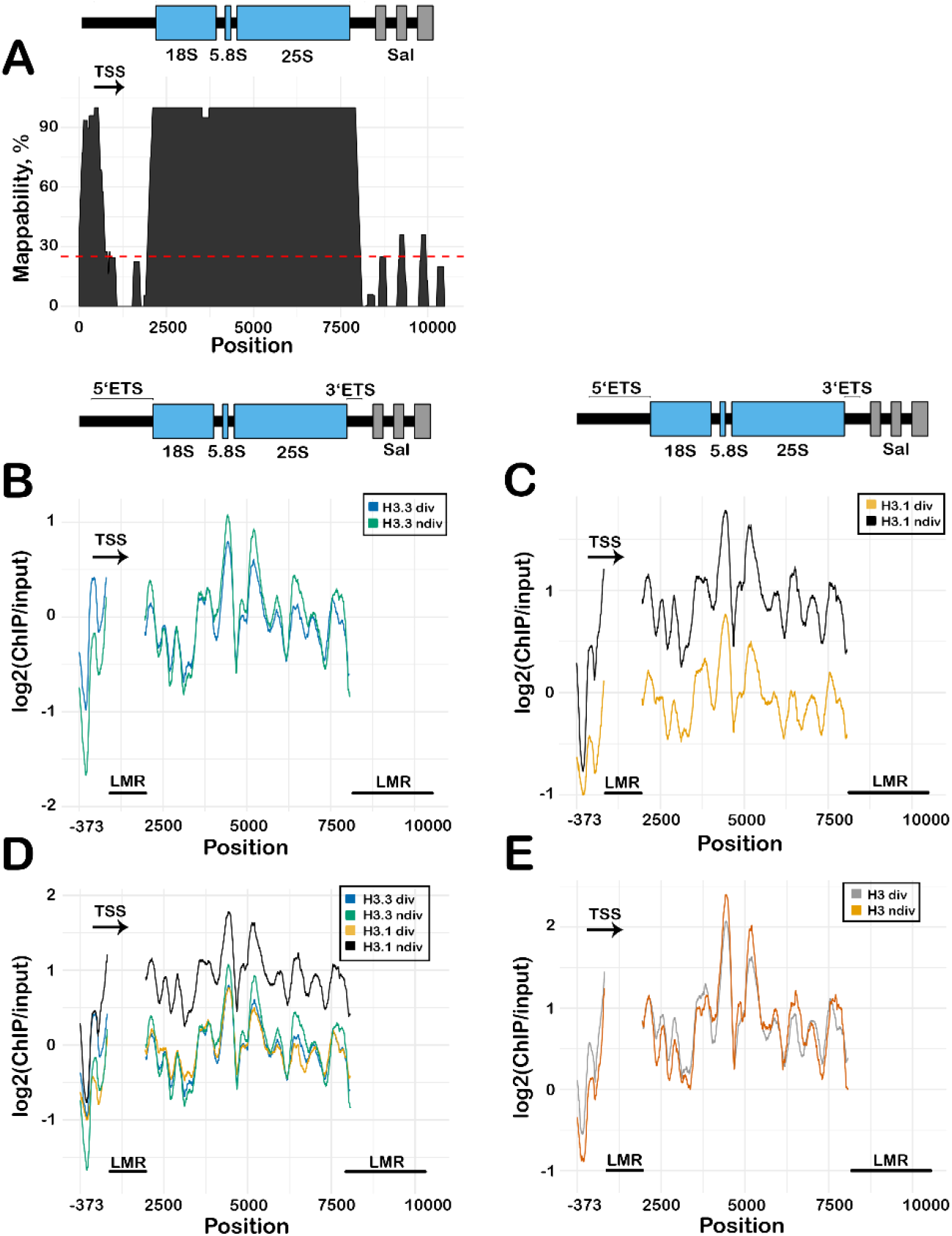
ChIP-seq read mappability and histone variant distribution in different tissues. (A) Schematic depiction of rDNA unit with graph showing the mappability of the rDNA sequence. Regions below the red line show less than 25% mappability and are considered unreliable for further analysis (labelled as LMR at further images). Distribution profiles (B) of H3.3 in dividing (H3.3 div) and non-dividing tissue (H3.3 ndiv); (C) H3.1 in dividing (H3.1 div) and non-dividing tissue (H3.1 ndiv); (D) H3.1 and H3.3 distribution profiles in dividing (H3.1 div, H3.3 div) and non-dividing tissue (H3.1ndiv, H3.3 ndiv); (E) H3 profile in dividing (H3 div) and non-dividing (H3 ndiv) tissue. Publicly available ChIP-seq datasets were used; ETS – external transcribed spacer; LMR – low mappability region; TSS – transcriptional start site.

**Fig. S5.**
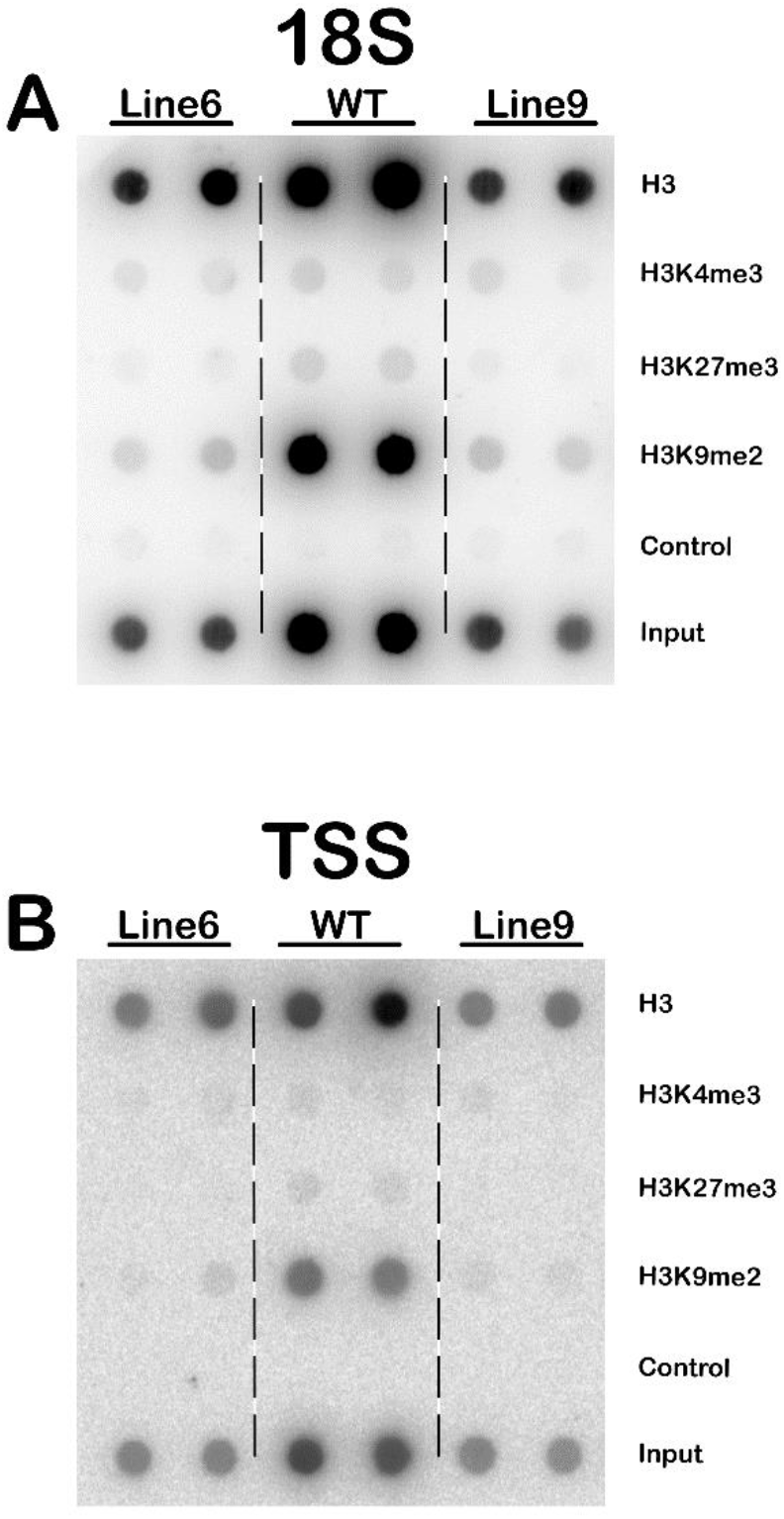
Analysis of histone modifications with ChIP and subsequent Dot-blot (long exposure) ChIP experiments with subsequent dot-blot signal detection showing levels of H3 histone and corresponding epigenetic marks at the (A) 18S and (B) TSS regions of a rDNA unit. Membranes were overexposed in order to visualize the weaker signals coming from some of epigenetic modifications.

**Fig. S6.**
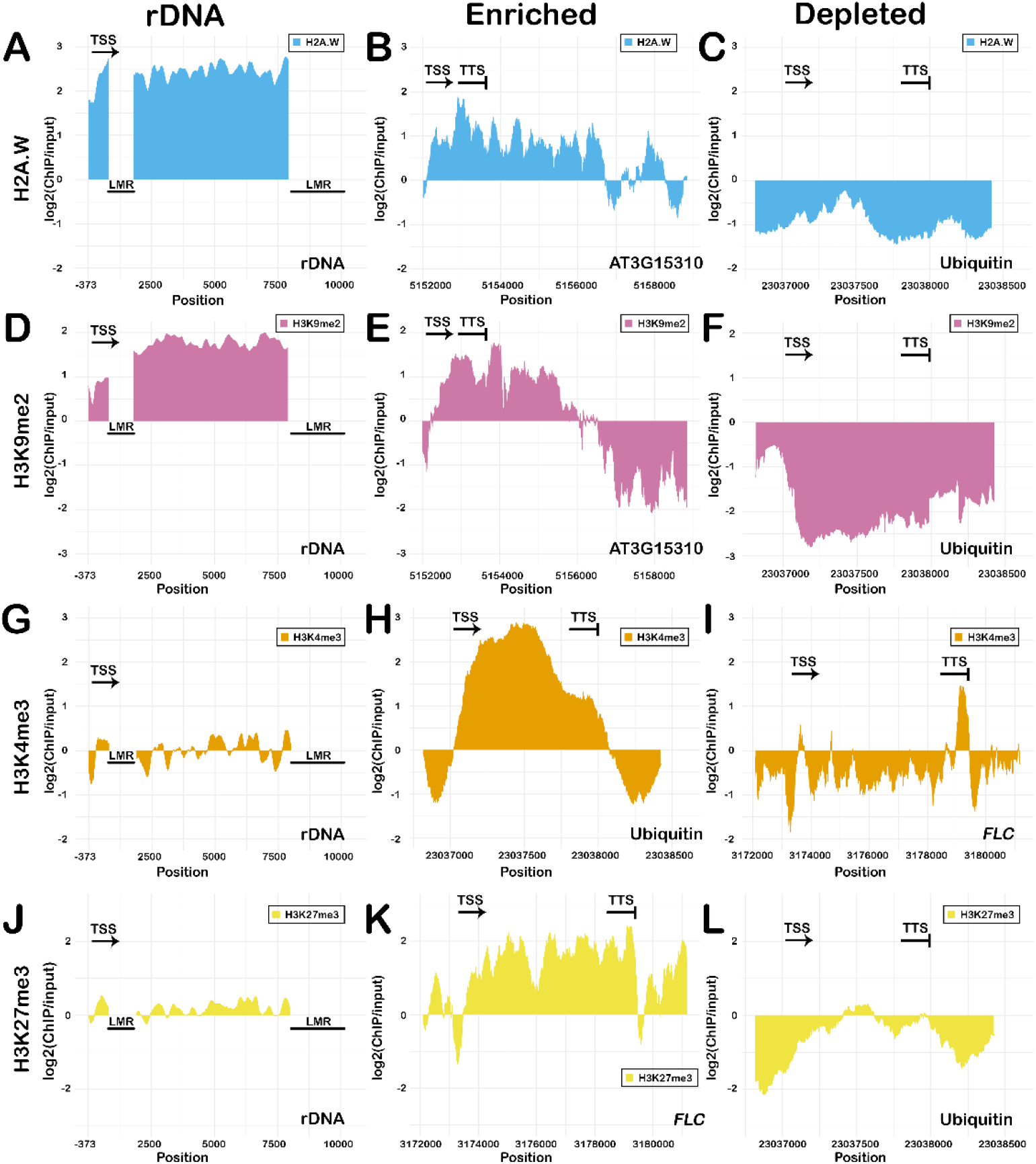
Comparative analysis ChIP-seq signal from epigenetic marks in rDNA and regions of enrichment or depletion. (A-C) H2AW and (D-F) H3K9me2 levels at (A and D) rDNA; (B and E) enriched region at AT3G15310 (transposable element); (C and F) depleted region at *UBIQUITIN 5* gene. G-I) H3K4me3 levels at (G) rDNA; (H) enriched region at *UBIQUITIN 5* gene; (I) depleted region at Flowering Locus C (*FLC*). (J-L) H3K27me3 levels at (J) rDNA; (K) enriched region at *FLC*; (L) depleted region at ubiquitinin5 gene. LMR – low mappability region, TSS – transcriptional start site, TTS – transcriptional termination site.

**Fig. S7.**
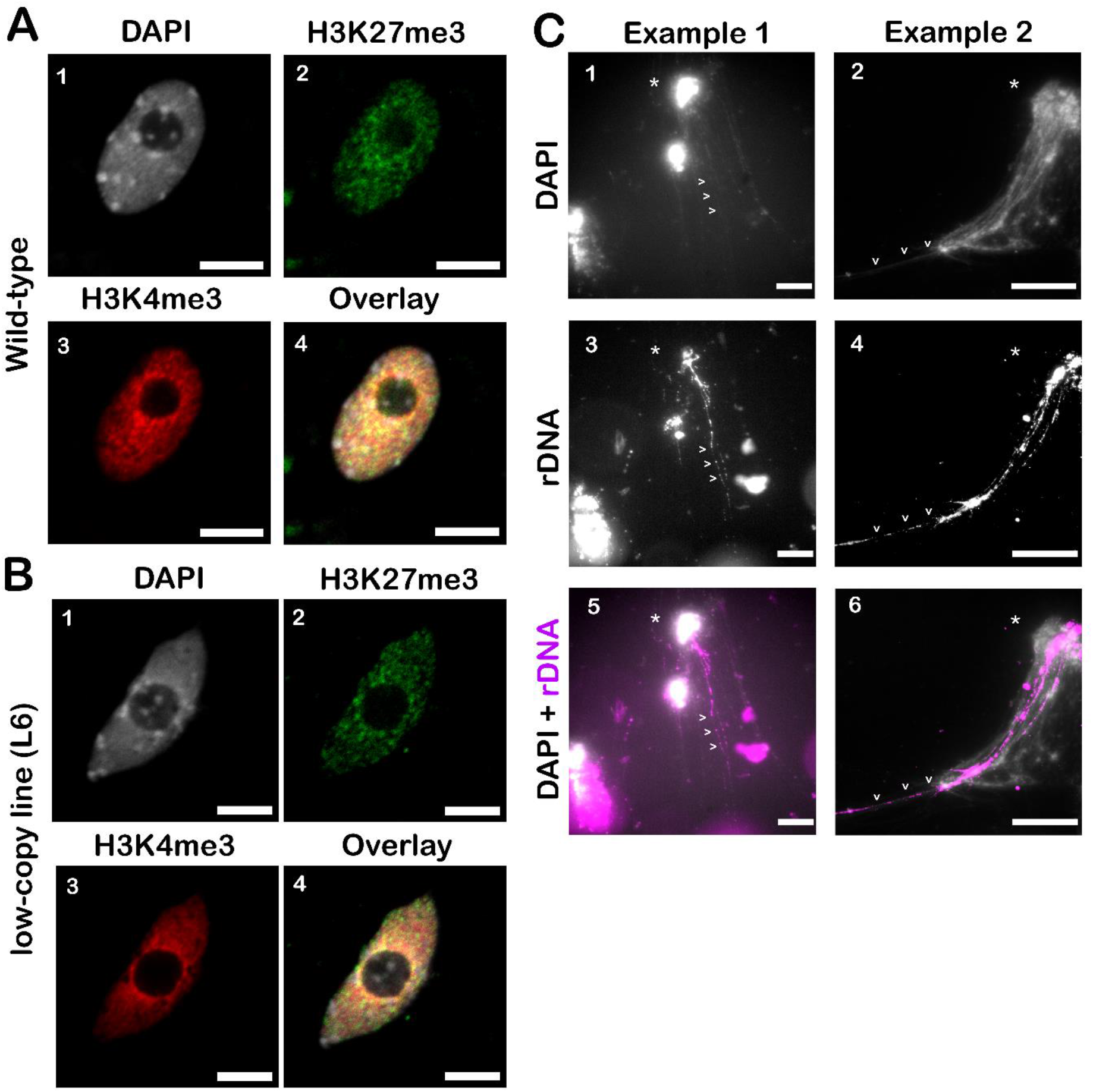
Visualization of histone marks in isolated nuclei and chromatin fiber FISH validation. (A) Distribution of histone marks (2) H3K27me3 and (3) H3K4me3 in nuclei of WT plants using confocal microscopy. (B) Analysis of histone marks (2) H3K27me3 and (3) H3K4me3 in low-copy plant line L6. (C) FISH on chromatin fibers targeting rDNA. (1-2) Chromatin fibers extending from partially lysed nuclei seen in DAPI. (3-6) Hybridisation of rDNA probe to some of some of the fibers extending from lysed nuclei. Partially lysed nuclei are marked by an asterisk. Chromatin fibers corresponding to rDNA clusters are indicated by arrowheads. Scale bar – 5 μm (A, B); 10 μm (C).

**Fig. S8.**
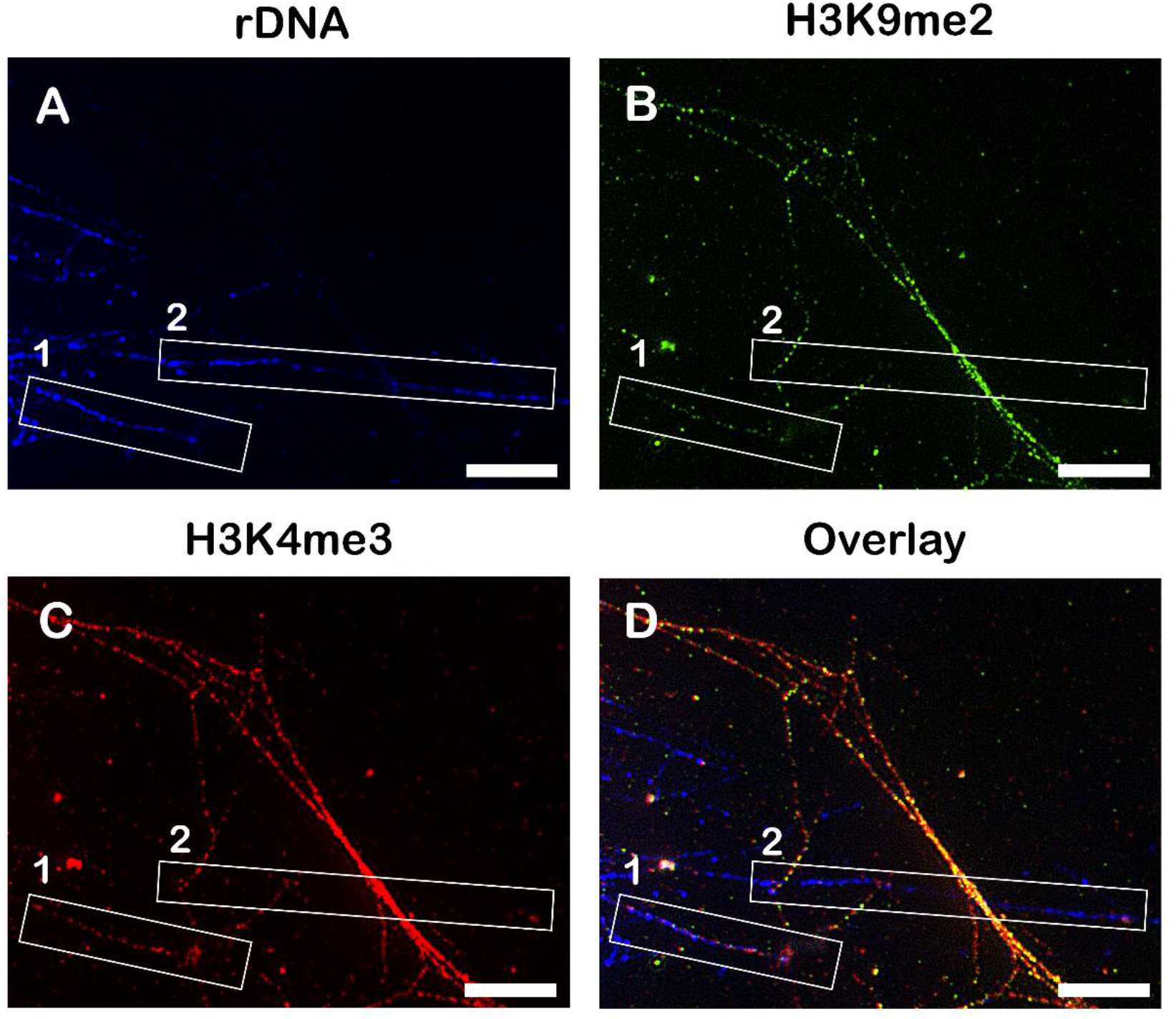
Detection of histone modifications in chromatin fibers (SIM-microscopy) (A) Signal from the rDNA probe marks several fibers in the field of view (FOV). Some are decorated with histone marks (rectangle 1) and some harbor nucleosome-free regions (rectangle 2). (B and C) Distribution of the repressive histone mark H3K9me2 (B) and activating histone mark H3K4me3 (C) on chromatin fibers. D) Overlay of the channels. Note that the absence of histone marks in rectangle 2 is not due to “histone stripping” during fiber extension, since other fibers in the FOV show strong signals from histone marks. Scale bar – 5 μm.

**Fig. S9.**
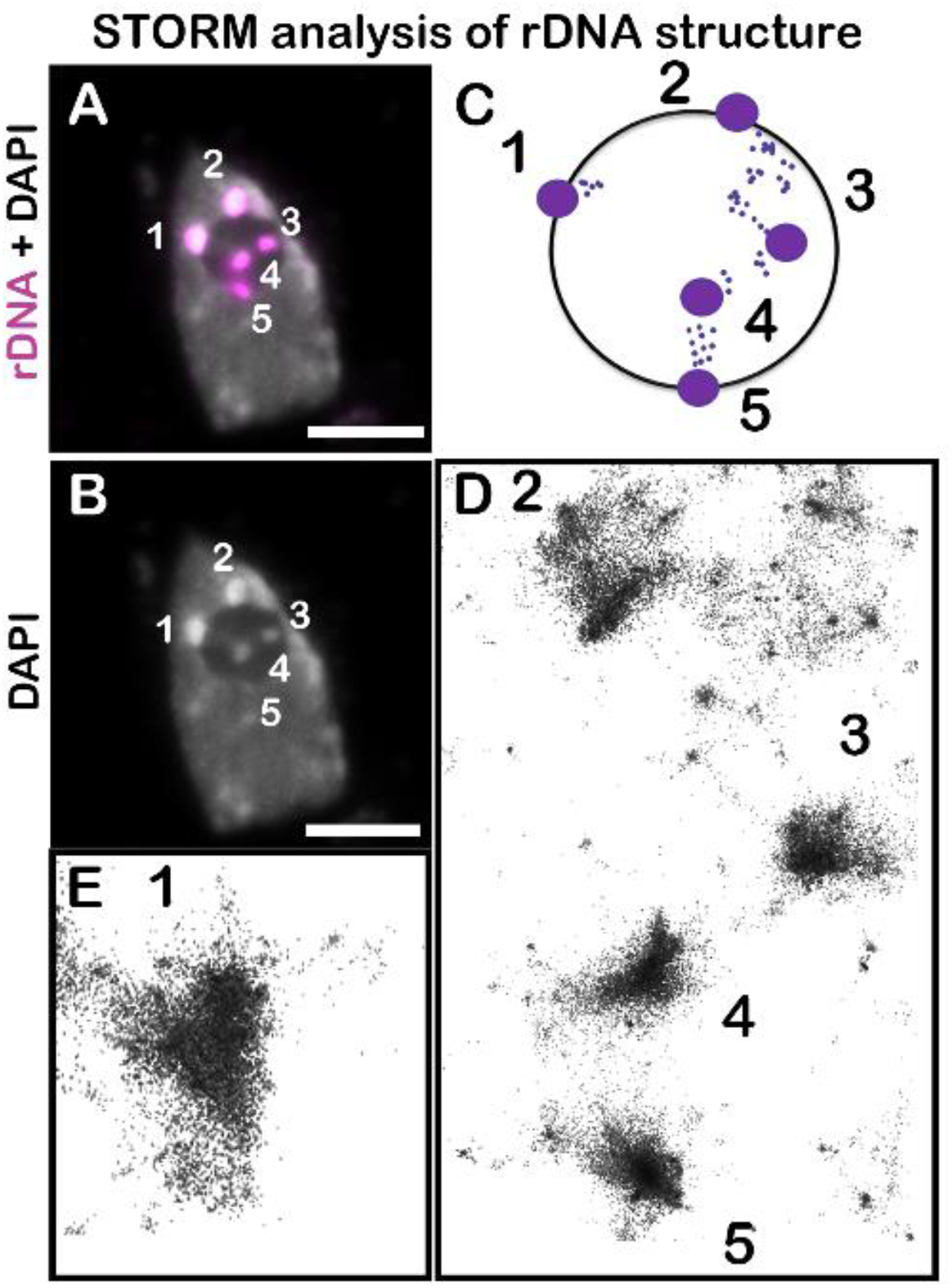
Ultrastructure of rDNA revealed by dSTORM. (A and B) Widefield image of a nucleus hybridized with a rDNA probe. Individual perinucleolar chromocenters (PNCs; 1,2,5) and nucleolar foci (NF; 3,4) are depicted schematically in (C). Some PNCs do not appear to extend into the nucleolus (E, 1), while rDNA signals between PNCs and NF can be seen in D (chromocenters 2-3; 4-5), suggesting possible connections. Scale bar – 5 μm.

